# Activity of tonically active neurons in the primate striatum reflects interaction between time processing and reward prediction

**DOI:** 10.1101/2022.10.07.511354

**Authors:** A.-C Martel, P. Apicella

**Author notes:** Author contributions :* ACM, Acquisition of data, Analysis and interpretation of data, Drafting or revising the article; PA, Experimental design, Acquisition of data, Analysis and interpretation of data, Drafting or revising the article.

## Abstract

The striatum and its dopaminergic input participate in temporal processing and numerous studies provide evidence that interactions between dopamine and acetylcholine are critical for striatal functioning. However, the role of local cholinergic innervation of the striatum in behaviors requiring precise timing has not been specifically investigated. Here, we recorded from presumed striatal cholinergic interneurons, identified as tonically active neurons (TANs), in two male rhesus monkeys performing self-initiated movements after specified learned time intervals have elapsed since a visual cue. We found that 38% of all recorded TANs responded to the cue with a pause in firing and the strength of these responses could be modulated according to the duration of the interval being timed and the accuracy of time estimates. By examining the TAN response to the reward itself and by recording from TANs during a Pavlovian procedure in which no action was required, we found evidence that TAN activity modulation may potentially reflect differences in the animal’s prediction of reward. Thus, besides their well-known role in predicting and detecting rewarding events, TANs may generate signals related to the processing of time. Our findings suggest a role of the local cholinergic circuitry in the representation of time within the striatum.

## Introduction

Convergent evidence indicates that the striatum, the main recipient of afferents to the basal ganglia, is a key component in brain networks involved in temporal processing. Apart from timing deficits observed in striatal-based disorders in humans, such as Parkinson’s disease (Pastor et al., 1992; Parker et al., 2013) and attention-deficit hyperactivity disorder (Rubia et al., 2009; Noreika et al., 2013), a large body of data supporting the role of the striatum and its dopaminergic input in the processing of time also comes from brain imaging studies in humans (Harrington et al., 2004; Coull et al., 2011) and lesion/inactivation studies in animals (Meck, 1996; 2006; Paton and Buonomano, 2018). In addition, electrophysiological studies in both rodents and monkeys trained on timing tasks have provided evidence of striatal activity related to the encoding of time at both single-neuron and population levels (Matell et al., 2003; Jin et al., 2009; Chiba et al., 2015; Gouvêa et al., 2015; Mello et al., 2015; Bakhurin et al., 2016; 2017; Wang et al., 2018; Zhou et al., 2020). Until now, hypotheses about the neuronal basis of striatal timing function have focused on the output neurons and the modulating influence of dopamine on temporal processing (Coull et al., 2011; Merchant et al., 2013; Paton and Buonomano, 2018). However, the striatum and its output pathways are strongly regulated by local circuits, including cholinergic interneurons, which exert a powerful influence on striatal network activity (Zhou et al., 2002). How the cholinergic innervation of the striatum, derived from interneurons, contributes to the processing of time remains unclear. On the other hand, there is considerable evidence that dopaminergic transmission interacts closely with the intrinsic striatal cholinergic system (Threlfell and Cragg, 2011; Cachope et al., 2012; Threlfell et al., 2012; Cai and Ford, 2018) and it is therefore possible that cholinergic signaling may complement DA transmission during timing behavior.

Although a link has been suggested between cholinergic activity and timing behavior based on lesion and pharmacological studies of cholinergic transmission in animals (Buhusi & Meck 2005; Meck 1996), none of them were specific enough to determine the contribution of the local cholinergic innervation of the striatum. Up to now, studies of behavioral effects of targeted disruption of striatal cholinergic transmission have documented disturbances in the ability to modify behavior in changing environments (Brown et al., 2010; Aoki et al. 2015), without specifically addressing a contribution to timing behavior.

Cholinergic interneurons of the striatum, presumed to be tonically active neurons (TANs) extracellularly recorded in behaving animals, are considered as detectors of motivationally salient stimuli whose timing cannot be precisely predicted. Indeed, in previous work, we showed that TANs are particularly sensitive to the expected time of rewarding stimuli (Sardo et al., 2000; Ravel et al., 2001), suggesting that they may have access to temporal prediction information (Apicella et al., 2006). However, no study to date has specifically investigated the contribution of the cholinergic TAN system to timing, under conditions which require precise temporal control of behavior.

Another open question is whether reward expectation may bias time-related signals in the striatum. As for dopaminergic neurons, TANs are highly sensitive to reward-predicting stimuli (Aosaki et al., 1994; Apicella et al., 1997) and some of them are involved in the encoding of reward prediction error (Apicella et al., 2009; 2011). The intertwining of reward prediction and temporal processing is the source of an inherent difficulty in disambiguating effects related to timing processes from those related to expectations about reward (Paton and Buonomano, 2018; Fung et al., 2021).

In the present study, we analyzed the activity of TANs during a time estimation task which requires monkeys to produce overt estimates of elapsed time. In this task, a visual cue is presented to the animals indicating that the reward is delivered for a self-paced movement executed at a specified time interval since the cue. We found that TANs displayed responses to the timing cue, some of them being scaled according to the duration of the interval to be timed and the accuracy of time estimates. Our results also indicate that the observed modulations in TAN activity may integrate reward prediction, in addition to the processing of time. Based on these findings, we suggest that the local cholinergic circuitry of the striatum could provide a neuronal mechanism that detect and process stimuli relevant for timing behavior and engage the striatal mechanisms more directly involved in the representation of time. A preliminary account of this work previously appeared in a short review (Martel and Apicella, 2021).

## Results

### Monkey’s ability to estimate time intervals

We examined neuronal activity while rhesus monkeys performed a time estimation task (TET) in which they made self-initiated movements based on time estimates (Figure 1A). In this task, animals were presented with a visual spatial cue that indicates reward can be obtained for making reaching movements towards targets after a specified time interval, either *short* (1.0, 1.3s) or *long* (2.0, 2.3s), has elapsed since the onset of the cue. A schematic representation of the sequence of events in the TET is given in Figure 1A.

**Figure 1.**
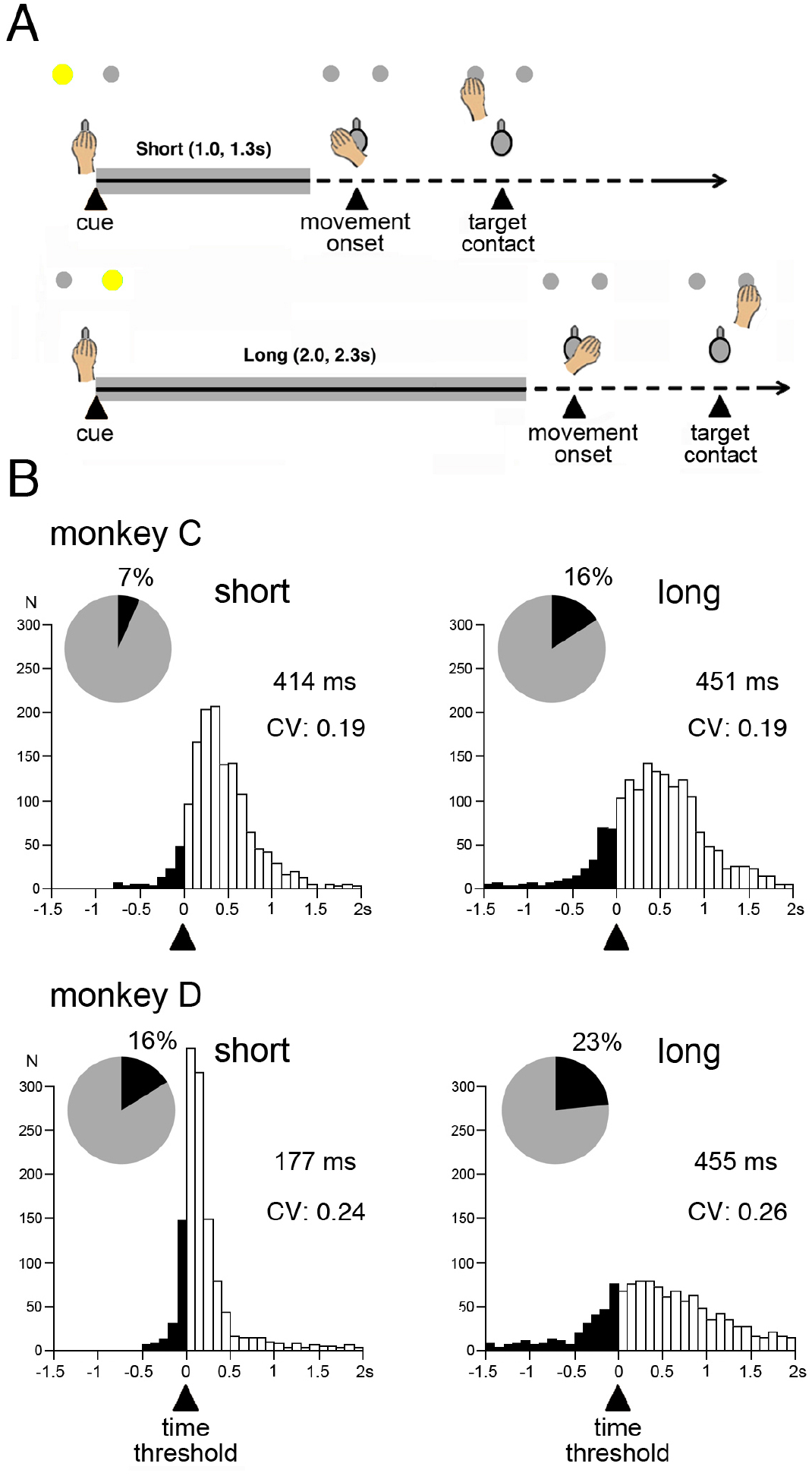
Time estimation task and timing performance. **A.** Temporal sequence of events in the task. Monkeys were required to initiate reaching movement after a specified time interval has elapsed. At the beginning of the trial, a visual spatial cue (yellow light), either on the left or the right side, indicated the duration of the interval *(Short* or *Long)* in the range of seconds, each time interval being associated with a particular location. Gray-shaded horizontal lines indicate the minimal waiting period before movement initiation assigned to each location of the cue (time threshold). Correctly timed movements were reinforced with fruit juice immediately after target contact. **B**. Timing performance. Distribution of movement onset times produced by monkeys in the task. CV, coefficient of variation (i.e., the ratio of the standard deviation to the mean).Values correspond to the mean onset time of movement after the criterion time. Black parts of the histograms correspond to movements which did not reach the criterion time (underestimation) and inset pie charts indicate the proportions of these incorrectly timed movements. Number of trials were 1370 and 1594 for monkey C and 1219 and 1146 for monkey D, for short and long intervals, respectively.

Distributions of movement onset times (MOTs) for each interval, indicate that the monkeys’ timing accuracy was lowered in both animals when the duration of the interval was longer, as reflected by a broader distribution around the time threshold for movement onset (Figure 1B). The relative constancy of the coefficients of variation in the range of intervals we used indicates that the extent of the MOT distribution was proportional to the length of the interval being timed, consistent with the scalar property of interval timing which refers to the tendency for response time variability to increase with interval duration (Gibbon, 1977). This supports the idea that both monkeys have learned to adjust the timing of their movements according to the location of the cue. We also quantified the timing accuracy of movements by calculating the percentage of trials in which animals correctly awaited the end of the minimum waiting period. Both monkeys responded after the criterion time in more than 80% of the trials, regardless of interval duration, reflecting their ability to adjust time estimates according to temporal information from cue. There was a mean 11.6% and 19.7% underestimation errors in monkeys C and D, respectively (Fig. 1B, insets), this difference being statistically significant (χ^2^ = 66.29, df = 1, p < 0.0001). In monkey D, the timing of movements was remarkably accurate for the short interval, the mean MOT being 177 ms after the time threshold, compared to 414 ms in monkey C. On the other hand, MOTs were relatively similar between the two animals for the long interval (451 ms and 455 ms in monkeys C and D, respectively). These interindividual differences may reflect different timing strategies. Namely monkey C could efficiently use the cue associated with each interval duration, leading to similar levels of timing accuracy in self-timed movements, whereas monkey D apparently used a strategy to optimize the estimated onset time of its movement specifically for the short interval.

### Sensitivity of TANs to the timing cue

We recorded the activity of 200 neurons (114 in monkey C, 86 in monkey D) electrophysiologically identified as tonically active neurons (TANs). Previous work in behaving monkeys has shown that TANs constitute a group of neurons that share characteristic spiking features (Figure 2-figure supplement 1) and display relatively homogeneous changes in activity related to the detection of motivationally salient stimuli (Apicella, 2017). In the present study, the predominant task-related modulation of TAN activity consisted in a short-lasting depression in activity (*pause*) after the presentation of the timing cue. This pause could be followed by a transient increase in activity (*rebound*) and, in some rare instances, preceded by a brief excitatory phase. The histological verification of recording sites in both monkeys showed that most of the recorded TANs were located in the dorsal putamen, at precommissural and postcommissural levels, as will be reported in detail later (Figure 7).

For each recorded TAN, we analyzed the responsiveness to the timing cue separately for short and long intervals. Over 38% of all TANs displayed statistically significant decreases in activity in response to the onset of the cue associated with the short and/or long interval (monkey C: 51/114, 45%; monkey D: 23/86, 27%), monkey C showing a larger fraction of responsive neurons than monkey D (χ^2^=6.80, df=1, p=0.009). Example responsive neurons are shown in Figure 2A. The first neuron (Fig. 2A, left) displayed a pause response to the cue for the short- and long-interval trials. In the second neuron (Fig. 2A, right), a pause response to the cue was detected for the short-interval trials, whereas a weak nonsignificant change occurred for the long-interval trials.

**Figure 2.**
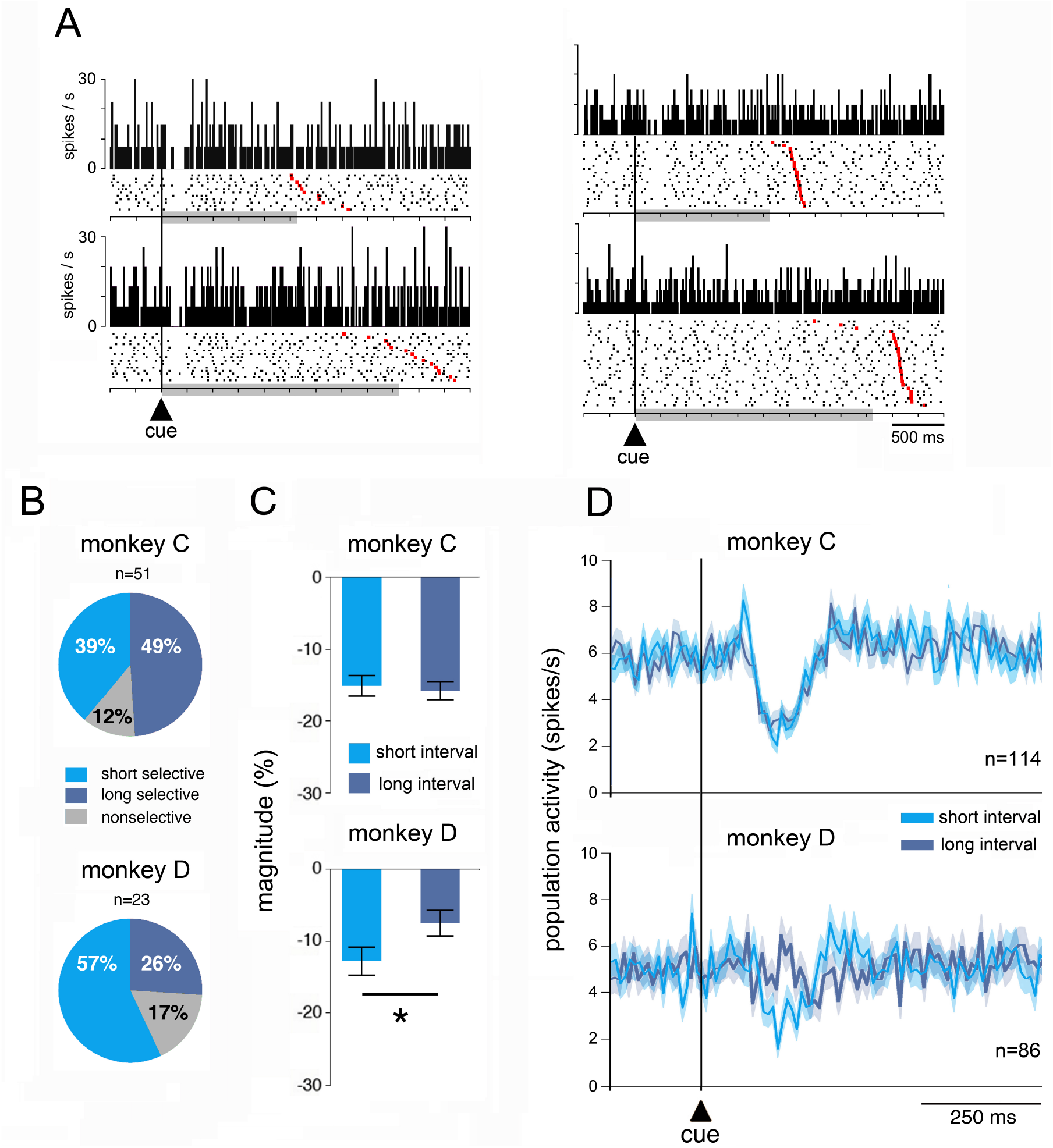
Sensitivity of TANs to the timing cue. **A.** Two example neurons responding to the cue. Left, nonselective response. Right, selective response. Each trial is displayed as a row of spikes (dots) aligned at cue onset, with perievent time histogram above each raster plot. Data are separated by interval duration and sorted by MOT. Red markers indicate the times of movement onset. Gray-shaded bars indicate the minimum waiting period before movement onset. **B.** Percentages of different response types evoked by the cue. n, number of responsive neurons. **C.** Boxplot representation of magnitudes of changes in TAN activity after the cue for short and long intervals. Data are indicated as decreases in percentage below baseline activity and expressed as means + SEM (* p < 0.05, Wilcoxon rank-sum test). **D.** Population activity of TANs aligned on the cue onset marked by the vertical line. The analysis included rewarded trials (i.e., correctly timed movements) and unrewarded trials (i.e., incorrectly timed movements). Colored curves represent mean activity of TANs separately for short and long interval trials calculated in nonoverlapping time bins of 10 ms. Shading indicates SEM. n, number of neurons included for population curves.

Of the 74 neurons responding to the cue, 10 (14%) responded regardless of the interval and 64 (86%) responded with only one interval, indicating that most TANs were able to discriminate between short- or long-interval cues. As illustrated in Figure 2B, in monkey C, there were no significant differences in the fraction of TANs showing selectivity for one interval or the other (short-preferring, n=20/51, 39%; long-preferring, n=25/51, 49%, χ^2^=0.99, df=1, p=0.318), whereas in monkey D the percentage of selective responses was significantly higher for the short interval than for the long interval (short-preferring, n=13/23, 57%; long-preferring, n=6/23, 26%, χ^2^=4.16, df=1, p=0.041). It therefore appears, in both animals, that most TANs showed selective responses to the cue depending on the interval duration, with monkey D showing an increased responsiveness of TANs to the cue associated with the short interval, possibly reflecting the higher accuracy of time estimates with this interval.

To analyze quantitatively whether the sensitivity of TANs to the timing cue differed according to interval duration, we rated the magnitude of changes in TAN activity during specified time windows which we selected on the basis of our analysis of latency and duration of individual neuronal responses for each monkey and each time interval (see Materials and Methods). We did this for each neuron of the entire sample of TANs tested in the TET. The results of this analysis are given in Fig. 2C. The magnitude of pauses in TAN activity varied insignificantly between short and long intervals in monkey C (Wilcoxon rank-sum test, z=0.415, p=0.677), whereas it was significantly higher for the short interval than for the long one in monkey D (Wilcoxon rank-sum test, z=2.241, p=0.025). We also examined the average activity from all neurons separately for the short and long intervals. As shown in Figure 2D, a clear population response after the cue onset was obvious for both intervals in monkey C, whereas a response occurred for the short interval in monkey D, with comparatively little sign of activity modulation for the long interval. In this animal, a preference for the short interval was therefore evident at the level of both the population and the individual neurons, reflecting the monkey’s tendency to be more efficient in using temporal information from short-interval cues. On the other hand, in monkey C, the magnitudes of the TAN response were not dependent of the interval duration, suggesting that this animal made use of temporal information to initiate actions in a more homogeneous way. This indicates that the different responsiveness of TANs in the two monkeys may potentially reflect differences in strategies for performing the timing task.

### Influence of timing accuracy on TAN responsiveness

We next examined whether TAN activity after cue onset may be related to the monkeys’ timing accuracy. To this end, we first compared movements initiated before reaching the time threshold (i.e., movements with negative MOTs) and movements that meet the time threshold (i.e., movements with positive MOTs), combining data from both intervals. As shown in Figure 3A, in monkey C, the magnitude of TAN pauses was stronger for incorrectly timed movements, compared with correctly timed movements (Wilcoxon rank-sum test, z=2.239, p=0.025), whereas differences in activity between correctly and incorrectly timed movements were not statistically different in monkey D (Wilcoxon rank-sum test, z=0.177, p=0.859) (Fig. 3A).

**Figure 3.**
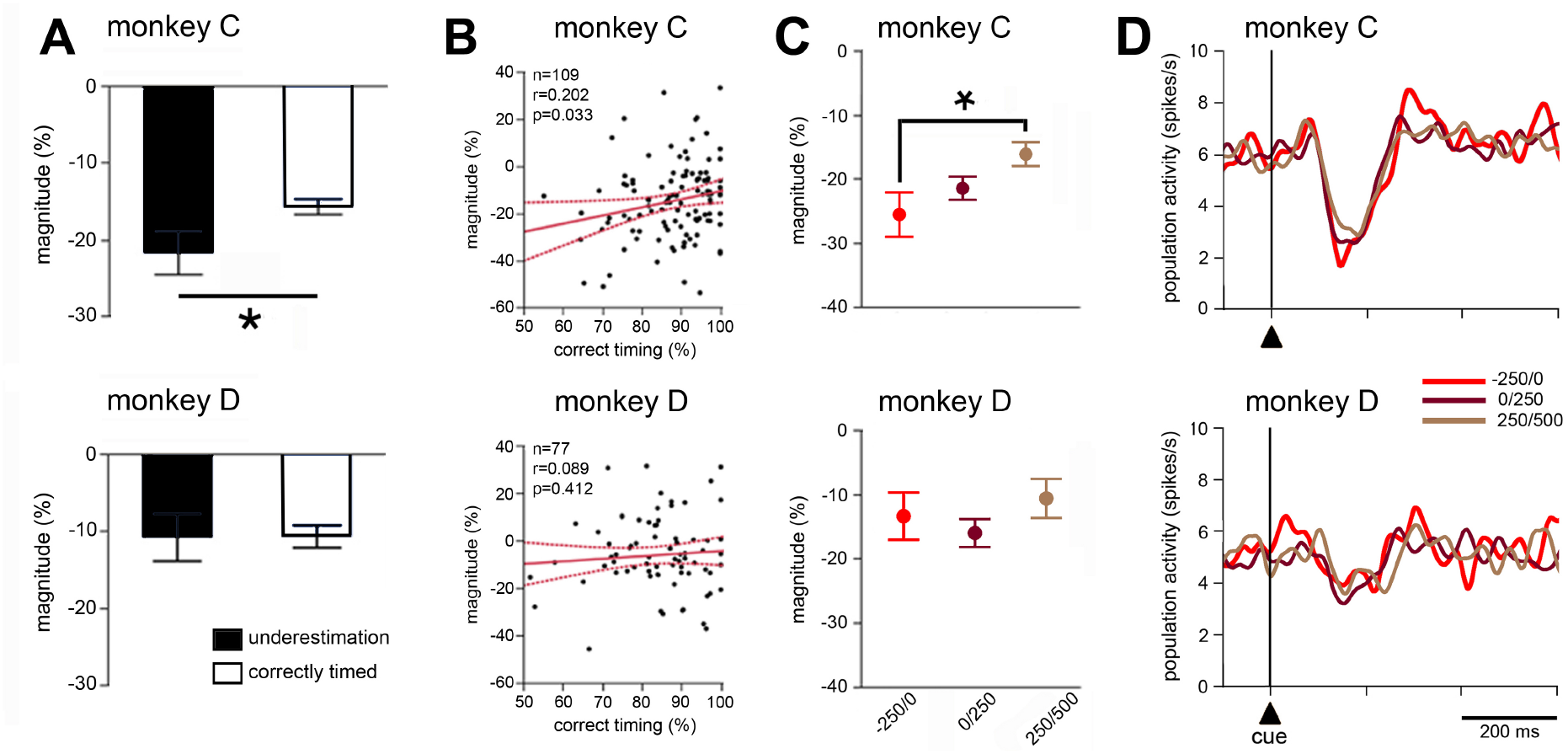
Sensitivity of TANs to timing accuracy **. A.** Comparison of magnitudes of TAN responses to the cue between correctly and incorrectly timed movements. Same conventions as in Fig. 2C. **B.** Correlations between magnitudes of TAN responses to the cue and levels of timing performance, for each block of trials in which a TAN was recorded. The solid lines (in red) indicate the fit of a linear regression and dashed lines indicate the 95% confidence interval from the regression. n, number of neurons. r, Pearson correlation coefficient. **C.** Influence of the proximity to the time threshold for movement onset. Magnitudes of TAN responses to the cue were computed separately from trials within different ranges of MOT (three consecutive 250 ms periods starting from 250 ms before to 500 ms after the time threshold), * *P*< 0.05. **D.** Population activity of TANs for trials in different MOT ranges. Same conventions as in Fig. 2D, except that curves are smoothed with a Gaussian filter (alpha=0.04).

To further assess a possible link between the TAN activity and timing performance, we performed a correlation analysis across neurons between the magnitude of TAN responses to the cue and the percentage of trials in which monkeys correctly reached the time threshold. As shown in Figure 3B, in monkey C, we found a statistically significant linear relationship between the_magnitude of TAN modulations and percentage of correctly timed movements (r=0.202, p=0.033), indicating that trials in which the magnitude of the TAN response to the cue was higher were followed by shorter production time intervals. In contrast, we did not found any significant correlation between the strength of the TAN response to the cue and timing accuracy in monkey D (r=0.089, p=0.412).

We then asked whether the strength of TAN modulation is dependent on proximity to the time threshold. To examine this, we sorted the trials by their MOT value and computed the magnitude of the pause response to the cue in three ranges of values around the time criterion (Fig. 3C). We found that the pause was significantly higher for the trials with the lower MOTs than for the trials with the higher MOTs in monkey C. This indicates that TAN response magnitude to the cue was influenced by the ability of monkey C at estimating the elapsed time before movement onset, with TAN modulations being stronger for movements initiated just before the time threshold (i.e., underestimation errors). Such a relationship between TAN response and timing accuracy was not observed for monkey D. However, when comparing across different MOT ranges in monkey C, there was no visible difference in the response of the population of neurons (Fig. 3D), indicating that individual TANs can discriminate better between the short- and long-interval cues than the average activity from the whole sample of neurons.

In summary, we showed, in one of our monkeys, that the strength of TAN response to the cue seemed to reflect the animal’s intention to move at a particular time, suggesting that TANs are involved in signaling motor initiation in a time-dependent manner. The interindividual difference in the sensitivity of TANs to timing performance emphasizes a possible link with the monkey’s strategy to determine the onset time of movements.

### Influence of reward prediction on TAN responsiveness

In the task employed here, the cue not only contains temporal information, but also predicts reward after a correctly timed movement. Because of the role TANs play in the detection of reward-predictive events, we should expect that prediction of the upcoming reward contributes to TAN sensitivity to the cue. Our behavioral data analysis showed that the proportion of correctly timed movements was higher in monkey C (88.4%) than in monkey D (80.3%), and this was paralleled by an increased TAN responsiveness to the cue in monkey C, suggesting that the increased ability of this animal at producing correctly timed movements favored predictable rewards and then enhanced the TAN responsiveness to the cue. To test this suggestion indirectly, we examined the effect of the reward itself on TAN activity. It was reasoned that a higher reward predictive value of the cue would lead to a decreased TAN responsiveness to reward, consistent with the inverse relationship between reward prediction and TAN responses to reward we have reported in previous studies (i.e., the more predictable the reward, the less strongly TANs respond to it). We then turned to the effects of reward on TAN activity using a similar time window analysis to that done for assessing the responsiveness of TANs to the cue. We found statistically significant pause responses to reward in 28 of 114 neurons (25%) and 31 of 86 neurons (36%) in monkeys C and D, respectively, the TANs in monkey C showing a tendency to lower, albeit not significant, responsiveness to reward than did TANs in monkey D (χ^2^=3.10, df=1, p=0.07). This indicates that reward prediction could potentially be incorporated in TAN responses to the timing cue in monkey C.

To further investigate sensitivity to reward for all TANs, we examined the magnitude of changes in activity during specified time windows adjusted to latency and duration of individual TAN responses to reward. As shown in Figure 4A, in monkey C, we found that TAN responses to reward were stronger for the long interval, compared to the short interval (Wilcoxon rank-sum test, z=16.30, p < 0.0001). This is in contrast with what we observed in monkey D, in which the effect of the delivery of reward on TAN activity was relatively mild and (did not differ) in magnitude between intervals (Wilcoxon rank-sum test, z=0.21, p=0.833). In addition, when comparing across neurons the magnitudes of activity changes following the cue and reward, we found an inverse relationship between cue and reward responses in monkey C (i.e., TANs that responded weakly to the cue responded strongly to reward) irrespective of interval duration (Fig. 4B) (Wilcoxon rank-sum test, z=0.69, p=0.487), providing support to the idea that differences in the responsiveness of TANs to the timing cue may partly result from differences in reward prediction. This effect was not present in monkey D (Wilcoxon rank-sum test, z=0.33, p =0.534). The above results suggested that changes in reward prediction, in at least one animal, could account for TAN sensitivity to the timing cue we observed.

**Figure 4.**
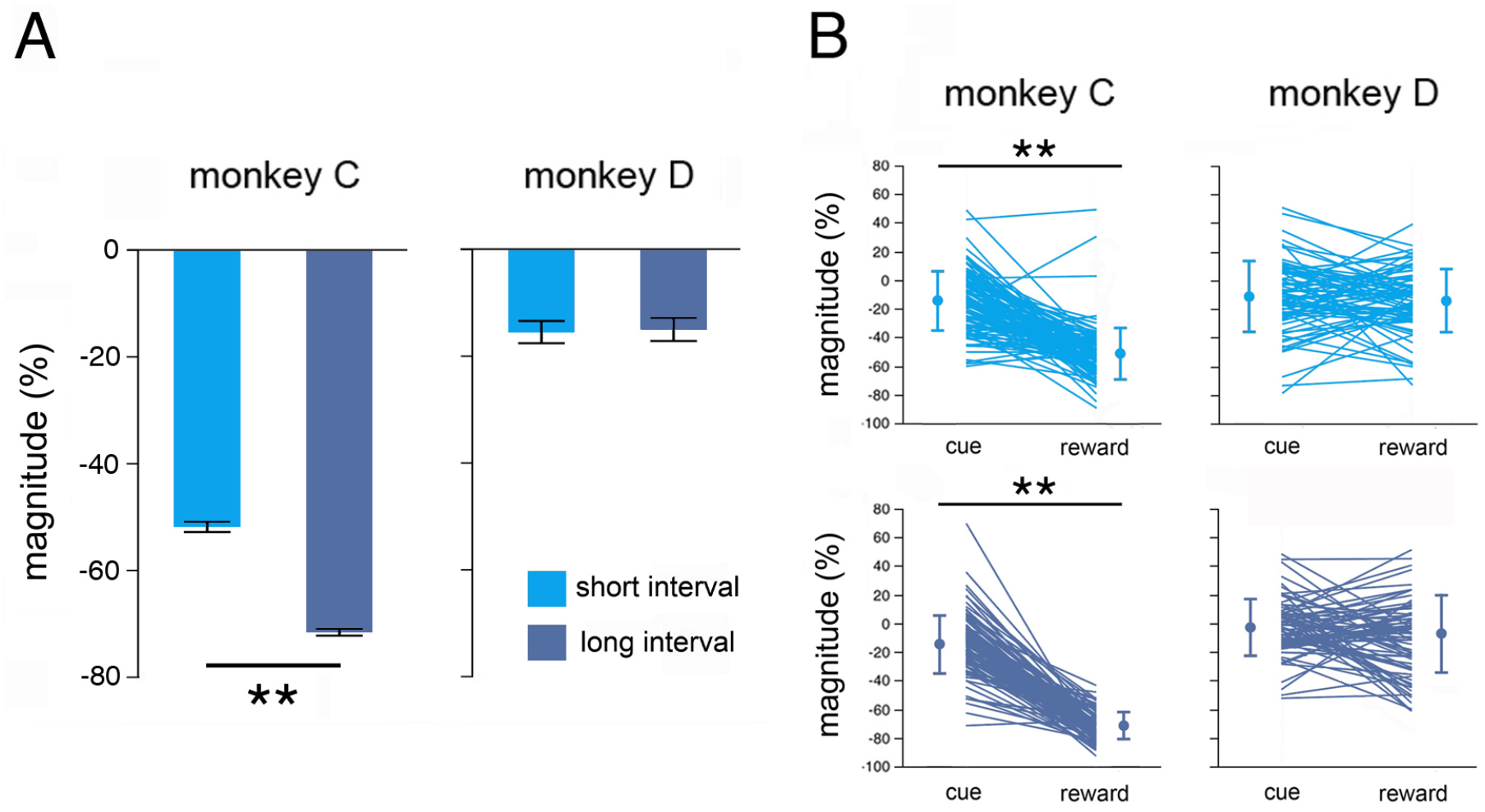
Sensitivity of TANs to reward **. A.** Boxplot representation of magnitude of changes in TAN activity after the delivery of reward separately for short and long intervals. Conventions are the same as in Fig. 2C, except that only rewarded trials were included. ** *P* < 0.01 (Wilcoxon signed-rank test). **B.** Changes in TAN activity after cue and reward separately for short and long intervals for each neuron recorded. Each line indicates the data of one neuron. Thick dots and error bars represent mean and SEM respectively. * *P* < 0.05, ** *P* < 0.01 (Wilcoxon signed-rank test).

In the TET, the movement was triggered by an internal decision process based on the estimation of elapsed time from the cue onset and there may be a possibility, in the absence of an external movement-triggering stimulus, that the movement itself serves as an event that predicts reward. We then focused on the influence of the movement onset on TAN activity. By analyzing TAN activity aligned on the initiation of movements in both monkeys, we showed that TANs, as a population, did not display activity modulation around the onset of movement, regardless of the interval duration (Figure 4-figure supplement 1).

### Influence of interval duration in the Pavlovian protocol

To assess more specifically the possible contribution of reward prediction in modulating TAN activity, we examined TAN responses when monkeys were engaged in a Pavlovian conditioning task (PCT) in which visual cues solely conveyed differences in the time elapsed until reward delivery, with no requirement for action. Such a condition allowed us to examine the influence of elapsed time while reward was invariably delivered on every trial. By using two time intervals in the same range as those in the TET and same relationships between cue location and interval duration, we found that both monkeys showed anticipatory licking movements prior to the delivery of reward indicating that the cue served as a predictor of reward (Fig. 5A).

**Figure 5.**
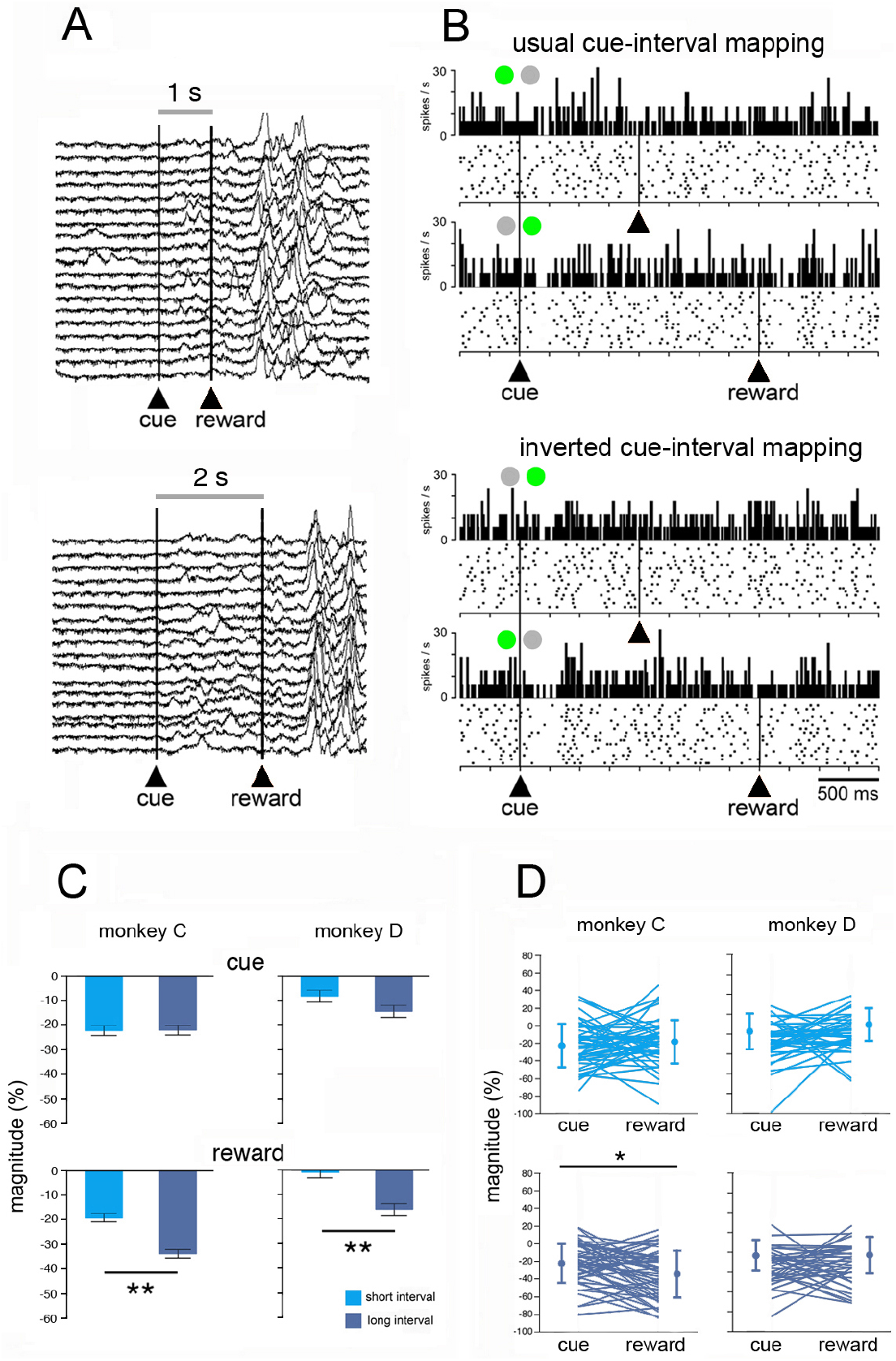
Effects of interval duration on TAN activity in the Pavlovian conditioning task (PCT). **A.** Licking movements after the onset of the reward predictive cue. For each interval duration are shown traces of lick records aligned on the cue onset. Licking started immediately after the presentation of the cue, indicating that animals have learned the reward predictive value of this signal. **B.** Raster plots of the activity of a single TAN recorded in the PCT. On this example, two blocks of trials were administered, corresponding to two cue-interval configurations. Rasters are separated by interval duration, with the upper two panels showing the TAN activity in the usual cue-interval configuration, and the lower two when the configuration has been inverted. The two circles at the top of each histogram indicate the left or right location of the cue (in green). Same conventions as in Fig. 2A. **C.** Magnitude of changes in TAN activity after cue and reward separately for short and long intervals. Same conventions as in Fig. 2C. **D.** Changes in TAN activity after cue and reward separately for short and long intervals for each neuron recorded. Same conventions as in Fig. 4B.

Among 90 TANs tested in the PCT, responses to the cue and reward were detected in 30 neurons (monkey C: 14/52, 27%; monkey D: 16/38, 42%) and 52 neurons (monkey C: 32/52, 62%; monkey D: 20/38, 53%), respectively. In contrast with what we observed in the TET, the proportion of TANs responding to the cue did not vary significantly between the two monkeys (χ^2^ = 2.27, df = 1, p = 0.131). Fig. 5B shows an example of a neuron sensitive to the interval duration in the PCT. In a first block of trials *(usual cue-interval mapping*), the neuron displayed a response to the cue which was stronger for long interval trials than for short interval trials. Then, the same neuron was tested in another block of trials (*inverted cue-interval mapping*) in which we reversed the assignments of the spatial cues for the short and long interval. As can be seen, the response selectivity was maintained, suggesting that the observed modulations in TAN activity did not appear to be related to the location of the cue. This neuron also responded selectively to reward delivered at the end of the long interval in both trial blocks. We examined the sensitivity to cue location in seven TANs by using standard and inverted cue-interval mappings and found that magnitudes of response to the cue did not depend on the spatial location of the cue (Figure 5-figure supplement 1).

As a next step, we undertook in the PCT a similar analysis to that done in the TET to assess quantitatively the effects of interval duration on changes in TAN activity within time windows determined for the cue and reward. As shown in Fig. 5C, the magnitude of TAN response to the cue failed to show any significant difference between time intervals, whereas a clear time-dependent modulation of reward response emerged in both monkeys, with stronger responses to long intervals compared to short intervals. This finding is consistent with results from our previous studies showing that a lack of prediction about the time of reward delivery has an enhancing effect on the TAN response to this event (Sardo et al., 2000; Ravel et al., 2001), which has been interpreted as reflecting a lowered monkey’s time-keeping ability with increased duration of the cue-reward interval. Contrary to what we observed in TET in monkey C, we did not find a clear relationship between cue and reward responses in PCT (Fig. 5D), with the exception of a weak, albeit significant, effect for the long interval (Wilcoxon rank-sum test, z=2.29, p=0.021). It therefore appears that interval duration exerts a more homogeneous effect on the activity of TANs in the PCT with the presence of a time-dependent effect on the reward response and the absence of relationship between cue and reward responses.

Lastly, we compared activity changes in a subset of 31 TANs recorded in both the TET and PCT. This comparison allowed us to differentiate the sensitivity of TANs to the timing cue when a similar underlying temporal representation in the seconds range was used either to produce an overt estimate of elapsed time (TET) or to simply predict reward after a time interval has elapsed (PCT). For some neurons, the strength of TAN modulation after the cue onset was different when passing from the TET to the PCT, while for other neurons cue responses remained constant in the two conditions. Examples of neurons responding to the cue are shown in Figure 6. The first neuron (Fig. 6, left) showed a weak response to the cue in the TET, while the cue response was more pronounced when the animal was shifted to the PCT, indicating that the response may vary with respect to changes in the learning context. As is apparent in the figure, the same neuron also responded to reward regardless of the condition, this response being greatest for the long interval. The second neuron (Fig. 6, right) showed a response to the cue that was clearly invariant to the condition. Although there was no modulation of activity following reward delivery in the TET, a response to reward was present only for the long-interval trials in the PCT. These findings indicate that some TANs showed different degrees of modulation of their activity upon variation of the task context in which timing cues were presented.

**Figure 6.**
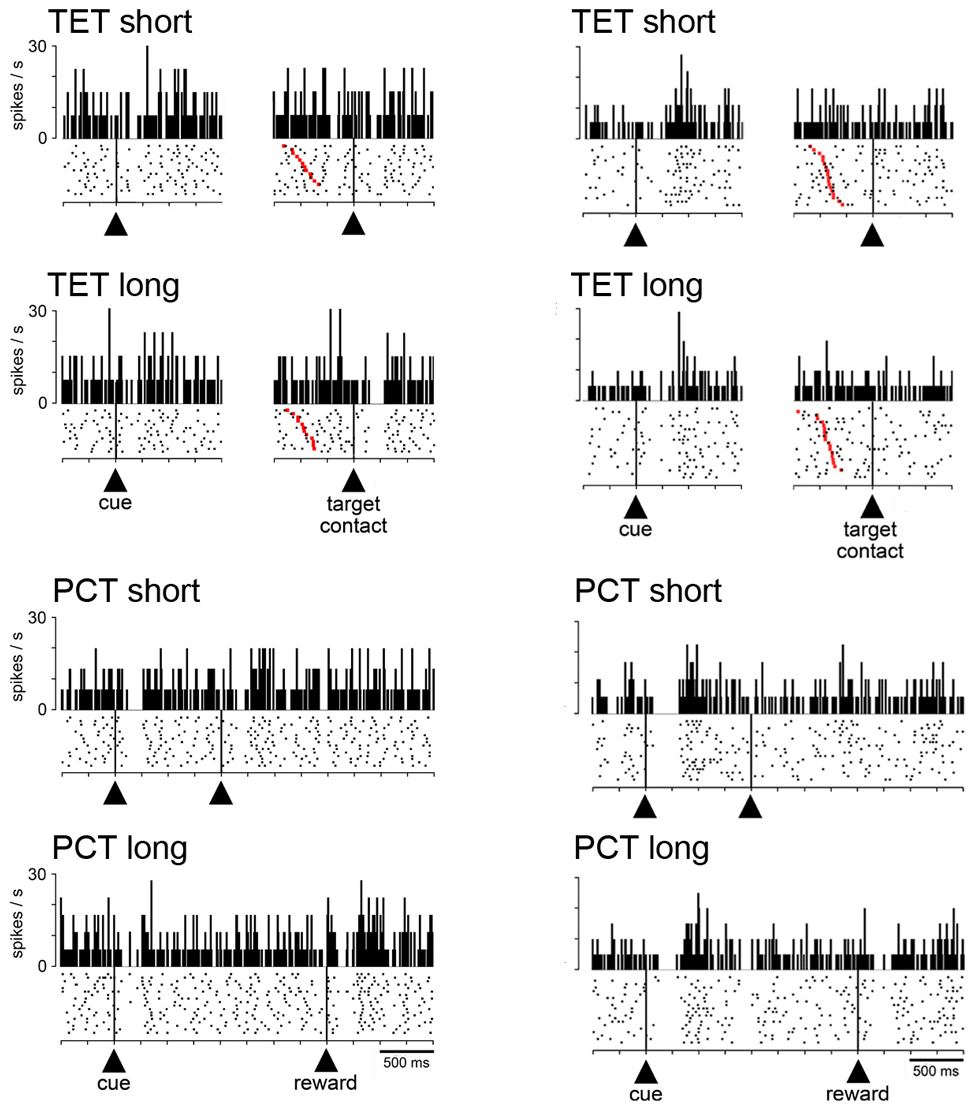
Influence of task condition on TAN responses to the cue and reward. Two example neurons tested in the two conditions. Rasters are separated by interval duration and task condition. Same conventions as in Fig. 2A, except that activity of each neuron is aligned separately to the cue onset and target contact in the TET. A few unrewarded trials are visible at the bottom of each raster aligned on reward delivery (absence of red marker) corresponding to monkey’s responses before the criterion time. TET, time estimation task; PCT, pavlovian conditioning task.

### Sensitivity of TANs to the timing cue among striatal regions

In primates, the posterior putamen is related to sensorimotor functions while dorsal and ventral parts of the anterior striatum are involved in associative and limbic functions respectively. In order to look for region-specific signals in TANs, the location of the recorded neurons was identified histologically in both monkeys. As illustrated in Figure 7A, TANs were mostly sampled from the dorsal parts of the precommissural and postcommissural putamen and were distributed across the same regions of the nucleus explored in the two animals. Only a small number of neurons (15 and 3 neurons in monkeys C and D, respectively) were recorded in the ventral striatum (i.e., the ventral part of the precommissural caudate nucleus and putamen). They were not assessed in detail because the sample size was too small for statistical analysis. We then compared the sensitivity of TANs to the timing cue in the two parts of the dorsal striatum (i.e., the motor and associative striatum). Responses to the cue were found in 37 neurons of the associative striatum (monkey C: 23/53, 43%; monkey D: 14/49, 29%) and 33 neurons of the motor striatum (monkey C: 22/46, 48%; monkey D: 11/34, 32%). Frequencies of cue responses did not vary significantly between the two regions (monkey C: χ^2^ = 0.195, df = 1, p = 0.658; monkey D: χ^2^ = 0.136, df = 1, p = 0.711), suggesting that response modulations related to the timing cue were not significantly different for the two regions of the striatum. Data from the two monkeys were also analyzed separately at the level of the population of TANs sampled in each dorsal striatal region, regardless of the responsiveness of individual neurons (Fig. 7B). The population responses to the timing cue in the motor and associative striatum were relatively weak in monkey D, compared with monkey C, and they did not appear to differentiate short- and long-interval trials.

**Figure 7.**
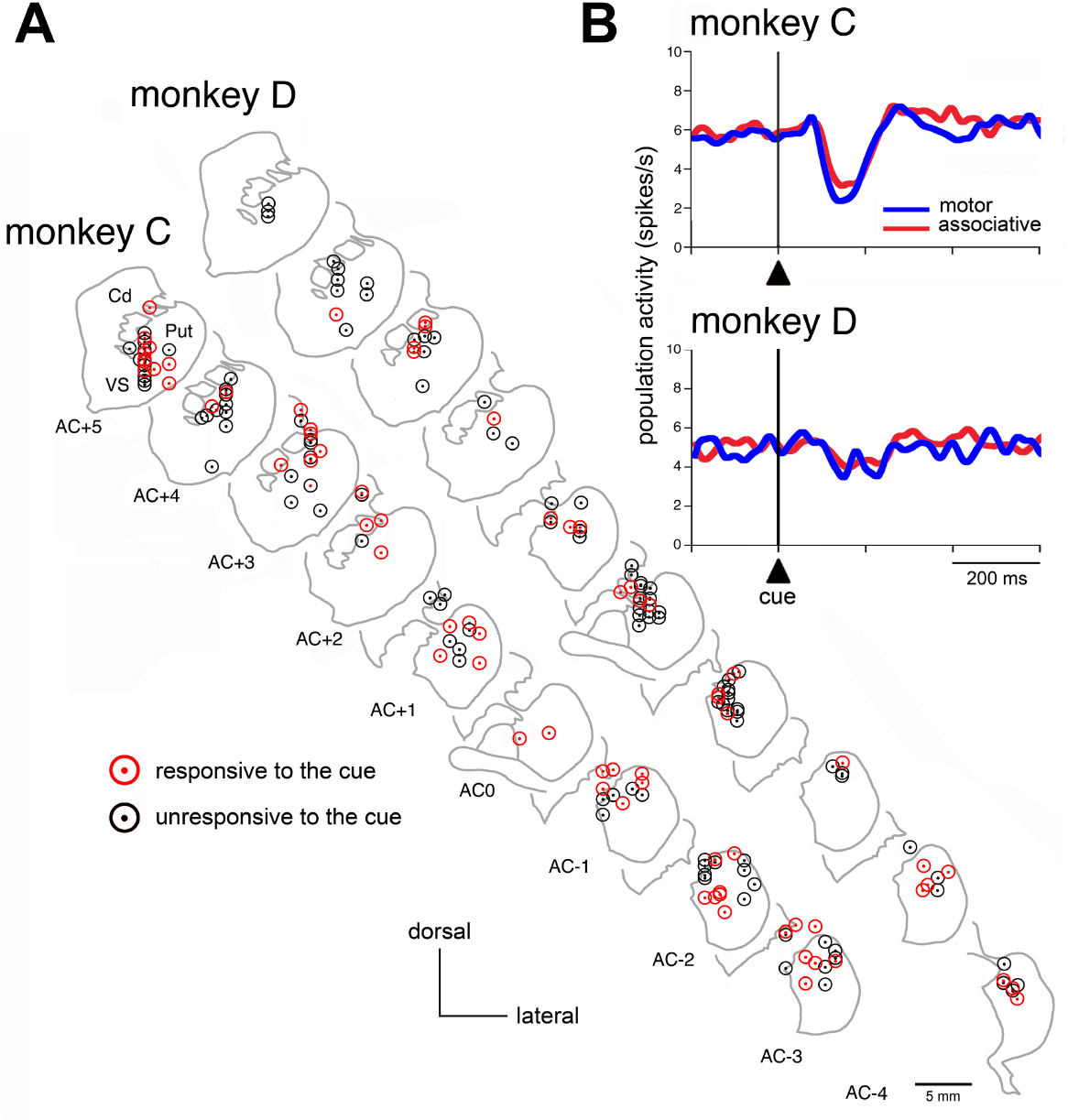
Sensitivity of TANs to the timing cue among striatal regions. **A.** Recording sites of TANs in the striatum of both monkeys. The location of all recorded TANs is plotted in the rostrocaudal direction on coronal sections from 5 mm anterior to 3-4 mm posterior to the anterior commissure (AC), with 1 mm intervals. Cd, caudate nucleus, Put, putamen, VS, ventral striatum. **B.** Comparison of population activities of TANs aligned on cue onset, grouped for short and long intervals, between motor and associative striatum. Same conventions as in Fig. 3D.

## Discussion

Previous experimental studies and theoretical models assessing the role of the striatum in the processing of temporal information have focused on the output neurons and did not outline any contribution of local circuit neurons which are critical regulators of the striatal network activity (Buhusi and Meck, 2005; Merchant et al., 2013). In previous work, we have documented that TANs, presumed cholinergic interneurons in the striatum, display strongest activity changes when rewarding stimuli are presented at unexpected times (Sardo et al., 2000; Ravel et al., 2001), suggesting that the integration of temporal information interacts with TAN signaling of reward-related events (Apicella et al., 2006). Here, we specifically addressed the role of TANs in temporal processing by analyzing their activity in monkeys performing self-paced movements based on time estimates in the range of seconds. Our results indicate that the responsiveness of TANs to the timing cue was influenced not only by temporal variables, but also by predictable rewards. This study provides the first account in behaving monkeys of modulation of TAN activity related to the animal’s ability to react in a temporally controlled manner, suggesting that the intrinsic striatal cholinergic system may contribute to timing function.

### TAN activity in relation to time estimation

To perform the TET, our monkeys were required to retrieve temporal information from visual cues, wait for a specified time interval, and then initiate a self-paced movement. Over 38 % of the TANs we recorded were responsive to the cue that marked the beginning of the time interval prior to movement initiation. Behavioral data confirm that the onset time of self-paced movements became more variable as interval duration increased, reflecting the increased variability in time estimates with the duration being timed, termed the scalar property of interval timing (Gibbon, 1977). Although both monkeys were trained extensively prior to recordings, our results revealed a difference in the degree of timing accuracy between the two animals, suggesting that they did not perform the task in the same way. On average, monkey D made more premature responses than monkey C and demonstrated an optimized timing performance for short-interval trials in terms of proximity to the time threshold. On the other hand, the timing strategy of monkey C did not change according to the interval duration, achieving approximately similar levels of timing accuracy for both intervals. Interestingly, these interindividual differences in timing performance were associated with differential responsiveness of TANs to the timing cue. The neurons in monkey C showed higher responsiveness to the timing cue (45%) than did neurons in monkey D (27%), suggesting that the timing cue was better taken into account, regardless of interval duration, for the former animal. Also, in monkey C, the strength of TAN modulation was similar for both short- and long-interval trials, whereas, in monkey D, TAN activity showed selectivity for the short interval compared to the long one, in terms of fraction of responsive neurons and the magnitude of responses. This may reflect, in this latter animal, the development of a spatial bias favoring the short-interval cue over the other. An intriguing result in monkey C was that TAN responses to the timing cue were stronger in trials with movements initiated before reaching the time threshold than in trials with correctly timed movements. We also found a relationship between the overall magnitude of TAN responses to the cue and the accuracy of timing (i.e., the stronger the neuronal modulation, the sooner was the self-timed movement). It therefore appears, at least for this animal, that the TAN response to the cue was specifically coupled to the onset time of the movement, providing support for the idea that TANs might participate in the adjustment of timed movements.

### Contribution of TANs to the encoding of time

The finding that TANs were responsive to the timing cue suggests a role for these presumed interneurons in initiating the timing process at the beginning of the interval to be timed, possibly through the shaping of sustained and/or sequential activity of striatal output neurons. Indeed, several recording studies in behaving animals have reported changes in the activity of output neurons of the striatum, at both single-neuron and population levels, that could convey information about how much time has elapsed (Matell et al., 2003; Jin et al., 2009; Chiba et al., 2015; Gouvêa et al., 2015; Mello et al., 2015; Bakhurin et al., 2016; 2017; Wang et al., 2018; Zhou et al., 2020). Our work supports the notion that the TAN response to the cue signals the detection of a stimulus associated with a learned time interval. This role for TANs may be quite similar to that proposed for midbrain dopaminergic neurons in a theoretical model of timing control, called the striatal beat frequency model, in which the dopaminergic response is considered as crucial for initiating the timing process (Matell and Meck, 2000; 2004; Buhusi and Meck, 2005; Meck, 2006). It is possible, based on known interactions between dopaminergic and cholinergic transmissions within the striatum (Threlfell and Cragg, 2011; Cachope et al., 2012; Threlfell et al., 2012; Cai and Ford, 2018), that signals arising from these two neuromodulatory systems provide striatal circuitry with a starting signal to keep track of elapsed time.

Apart from the characteristic brief firing rate changes observed at the level of individual neurons, TANs are also known to exhibit synchronous activity during behavior (Raz et al., 1996; Kimura et al., 2003; Morris et al., 2004) and one cannot exclude that synchronized firing among TANs that was not detected by our analyses may also participate in the encoding of time. Recent work using two-photon calcium imaging and population recording with cell-type-specific labeling in behaving mice has reported modulations of the cholinergic tone during spontaneous behavior which may impact striatal output dynamics (Howe et al., 2019). It is therefore conceivable that other encoding mechanisms across the cholinergic population may represent temporal information.

Our findings show that TANs responsive to the timing cue were distributed across the anterior caudate and putamen, a region assumed to serve cognitive-related functions, and the posterior putamen, which is important for motor control. Previous electrophysiological studies have reported that output neurons of the striatum, which make up the vast majority of neurons in this structure, convey temporal information both in motor and non-motor striatal regions in rodents (Matell et al. 2003; Mello et al., 2015; Gouvêa et al. 2015; Bakhurin et al., 2016; 2017; Zhou et al. 2020) and monkeys (Jin et al., 2009; Chiba et al., 2015; Wang et al., 2018). Although these studies have focused on dorsal striatal activity, at least one of them has reported that output neurons located in the ventral striatum, a region related to affective and motivational states, are involved in encoding time (Bakhurin et al., 2017). In our study, the responsiveness of TANs in the most ventral parts of the anterior striatum have not been assessed in detail, because the data set was small, and further recordings from this region are needed to draw conclusions about the potential contribution of ventral striatal TANs to temporal processing.

### Reward prediction and timing processes

In our task, the cue marks the beginning of the interval to be timed and indicates that reward is delivered for responding after a specified duration has elapsed. It therefore remains unclear whether TAN responses to the cue reflect time processing, as distinct from expectations about reward. It is increasingly recognized that timing and motivation processes are concurrently active during tasks used to elicit timing behavior in animals and humans (Fung et al., 2021). For example, a link between time estimation and reward expectation has been demonstrated for midbrain dopaminergic neurons in mice performing a temporal discrimination task (Soares et al., 2016). In the present study, we have assessed the sensitivity of TANs to reward prediction via TAN responses to the reward itself, since it is well established that fully predicted reward elicits little or no TAN response relative to reward delivered in an unpredictable way (Sardo et al., 2000; Ravel et al., 2001). As we found in monkey C, the magnitude of the cue response was negatively correlated with the magnitude of reward response across TANs, a stronger response to the cue being paralleled by a weaker response to reward irrespective of interval duration. This supports the notion, at least in this animal, that reward prediction could potentially be incorporated in TAN responses to the timing cue. In contrast, such a correlation was not observed in monkey D indicating little or no contribution of reward-predictive value of the cue, likely because this animal was less successful in obtaining rewards in the timing task, compared to monkey C. It is also noticeable that the overall magnitude of TAN responses to reward in monkey D was relatively weak, perhaps reflecting a lower level of motivation to perform the task.

In an attempt to assess more specifically the influence of reward prediction on the sensitivity of TANs to the timing cue, we used the PCT in which timing relies on expectation of when reward will be delivered. In this condition, differences in the magnitude of the TAN response to reward between short- and long-interval trials were consistent in both monkeys, response magnitude being higher when the timing of reward became less precise with longer intervals. Based on these findings, we suggest that TANs show a clear reward prediction effect in the PCT. On the other hand, the magnitude of the TAN response to the cue did not show a consistent relationship with the magnitude of the reward response, indicating that the modulation of the response of TANs to the reward-predicting stimulus was less marked than that following reward delivery, as previously observed in a study manipulating reward probability during a Pavlovian task (Apicella et al., 2009).

Although a clear-cut dissociation between temporal and motivational variables is difficult to make in the TET, our analysis of TAN responses to reward indicates that reward prediction should be considered as a possible confounding effect. This is in line with studies showing that the motivational state may affect time processing abilities (Gable and Poole, 2012; Avlar et al., 2015; Balci, 2014; Kirkpatrick, 2014; Daniels and Sanabria, 2017). The few neuroimaging studies in humans that have attempted to dissociate reward and timing processes, have reported activations in striatal and prefrontal cortical areas that may reflect the integration of time perception and reward expectation (Tomasi et al., 2015; Apaydin et al., 2018). In addition, theoretical models incorporating time as a crucial aspect of reward processing also emphasize the importance of motivational variables on timing processes (Daw et al., 2006; Galtress et al., 2012; Gershman et al., 2014; Petter et al., 2018; Mikhael and Gershman, 2019). Further work using behavioral procedures that will allow independent variation of temporal and reward parameters is required to clarify whether TANs participate in time processing independently of reward prediction.

### Context detection and timing processes

By focusing on a subset of TANs recorded in the TET and PCT, we sought to directly compare the efficiency of specific behavioral contexts for eliciting TAN responses to a cue which serves to either estimate a duration or simply predict the expected time of reward using short and long intervals in the same range. Results suggest that some TANs can adapt their responsiveness by taking into account the condition in which the timing cue is presented. This differential sensitivity to the timing cue indicates that the encoded information is likely to change with the particular learning context in which the cue was experienced, suggesting that TANs do participate in temporal processing in a context-dependent manner. This is in agreement with the notion that the local cholinergic circuitry of the striatum could play a role in the recognition of a learned context for stimulus detection and action generation (Apicella, 2007; Stocco, 2012; Bradfield et al., 2013). Single neuron recordings in behaving animals have reported in many circumstances an influence of context on TAN responses to salient events (Shimo and Hikosaka, 2001; Yamada et al., 2004; Lee et al., 2006; Apicella et al., 2011; Stalnaker et al., 2016).

Although dopaminergic neurons and cholinergic TANs are able to transmit signals that convey temporal information (Martel and Apicella, 2021), their contribution to the processing of time may not be equivalent. From an anatomical viewpoint, dopaminergic neurons send information to different brain regions, including the striatum and frontal cortex, thus exerting a widespread influence on multiple cortical and subcortical targets involved in the production of reward-guided behaviors, whereas the influence of the TAN signal is confined within the striatal circuitry. One can speculate that this local signaling system relates to contextual information that may impact behavior. The fact that the TAN response is modulated by multiple factors, including possible individual strategies for initiating self-timed movements, is consistent with the notion that the information encoding capacity of the TAN system might specify the context upon which timing is applied.

### Limitations of the study

In the present study, monkeys produced either a short or a long time interval by making spatially directed movements. An obvious limitation of this design is that the cue not only provided a temporal reference for self-initiated movements, but also contained information about which action to select, each time interval being associated with a movement to a right or left target. Therefore, we cannot rule out an influence of the visuospatial nature of the timing task on the TAN responses to the cue we observed. Although previous studies have shown that TANs may display changes in activity in response to visuospatial aspects of task performance (Kimura, 1992; Shimo and Hikosaka, 2001; Ravel et al., 2006), it is generally acknowledged that TAN activity does not have a consistent relationship with the production of a specific action (but see Lee et al., 2006 for the observation of TAN modulations related to self-paced movements). In addition, TANs have been proposed to drive attentional shifts required for stimulus detection (Matsumoto et al., 2001; Ding et al., 2010; Schulz and Reynolds, 2013), suggesting that differences in the monkey’s allocation of attention across the two cue locations may bring about differences in the TAN response. However, in the Pavlovian protocol, we reversed the relationship between the location of the timing cue and the interval duration and did not find compelling evidence that a spatial effect may account for observed changes in TAN activity. Additional work is needed to disambiguate possible effects related to time processing from those related to visuospatial attentional demands.

To conclude, our results support a view that TANs appear to act like detectors of stimuli relevant for initiating a timing process in the seconds range, providing evidence that the information encoding capacity of TANs extends to cognitive functions, including the representation of time. Although identifying the involvement of TANs in timing behavior remains challenging due to the difficulty to distinguish signals specific to temporal processing from those related to other confounding processes, changes in TAN activity we observed likely reflect a combination of time perception and reward prediction. An important question for future studies will be to clarify how the coordination between the TAN system and striatal output pathways, under the control of dopamine, participates in striatal timing function. Until now, there is no experimental evidence from inactivation studies in animals that a selective alteration in intrinsic striatal cholinergic innervation disrupts timing behavior. However, human studies have pointed to an impaired temporal control of motor behavior in patients with Tourette syndrome (Vicario et al., 2010; Graziola et al., 2020; Schuller et al., 2021) in which a loss of striatal cholinergic interneurons has been reported (Kataoka et al., 2010). There is also strong evidence that striatal dopamine-depleted states are associated with disturbances in timing behavior in both animals and humans (Buhusi and Meck, 2005; Merchant et al., 2013), and it is conceivable that impaired local interactions between dopaminergic and cholinergic transmissions in the striatum are central to the expression of time processing abnormalities. Future work will be necessary to fully understand the integration of cholinergic TANs and dopaminergic input into striatal circuits underlying the encoding of time.

## Materials and methods

### Animals and recording procedures

Experiments were conducted on two adult male rhesus monkeys (*Macaca mulatta*), C and D, weighing 11 and 7 kg, respectively, kept in pairs in their home cage. This study was performed in accordance with the principles of the European Union Directive 2010/63/EU on the protection of animals used for scientific purposes and all procedures were approved by the Ethics Committee of the Institut de Neurosciences de la Timone (protocol #3057-2015120809435586). Prior to the experiments, each monkey was implanted with a head-holding device and recording chamber (25×35 mm) under general gas anesthesia (sevoflurane 2.5%) and aseptic conditions. The center of the chamber was stereotaxically directed to the anterior commissure (AC) based on the atlas of Paxinos et al. (1988). The animals received antibiotics and analgesics for a period of 5 days after the surgery.

Recordings were obtained with glass-coated tungsten electrodes (impedance: 2–3 MΩ) passed inside a guide tube (0.6 mm outer diameter) and lowered to the striatum with a manual hydraulic microdrive (MO-95; Narishige, Tokyo, Japan). Electrode tracks were made vertically into regions of the striatum anterior and posterior to the AC, mostly in the putamen. Few penetrations were continued through the precommissural caudate nucleus and putamen to reach the ventral striatum.

Neuronal activity was amplified (x5000), bandpass-filtered (0.3–1.5 kHz), and spike sorting was performed on-line using a window discriminator (NeuroLog; Digitimer, Hertfordshire, UK). Continuous monitoring of the spike waveform on a digital oscilloscope allowed us to check the isolation quality of the recorded neurons. Recordings took place in the left striatum and all reach movements were made with the arm contralateral to the recorded hemisphere. In line with many previous single-neuron recording studies in the striatum of behaving monkeys, TANs were electrophysiologically identified by waveform shape, firing rate, and response to motivationally salient stimuli consisting of a transient decrease in activity (*pause*). A computer controlled the behavioral task and data acquisition using a custom-made software developed by E. Legallet under LabVIEW (National Instruments).

### Behavioral task

Training and recording sessions took place in a setup similar to that described previously (Marche and Apicella, 2017). Monkeys were seated in a restraining box facing a vertical panel containing two metal knobs (diameter, 10 mm) serving as movement targets, positioned 18 cm apart at the animal’s eye level, and two light-emitting diodes (LEDs), one above each target. A resting bar was mounted in the lower part of the panel and a tube was positioned directly in front of the monkey’s mouth for dispensing drops of fruit juice (0.3 ml) as a reward. Animals were trained in a *time estimation task* (TET) in which one of the two LEDs was lit as a cue indicating both the target of movement and the minimum waiting period before initiating a reaching movement, the left and right locations being associated with *short* and *long* intervals, respectively. A schematic representation of the sequence of events in the TET is given in Figure 1A. The trial began when the monkey kept its hand on the bar. After 1 s, a LED was illuminated for 0.5 s. This cue marked the beginning of the to-be-timed interval, referred to as the time threshold, and its left or right location indicated that the minimum waiting period before movement onset was short or long, respectively (monkey D: 1.0 and 2.0 s; monkey C: 1.3 and 2.3 s from cue onset). The onset of movement after the minimum waiting period has elapsed was limited to 2 s. The cue location was pseudorandomly selected from trial to trial. Trials lasted 6 s, so that the overall temporal structure of the task remained the same. Because of this fixed trial duration, the monkey had no possibility to predict the timing of the cue accurately while keeping the hand on the bar.

Only monkeys’ reaching movements made after the specified time interval had elapsed were rewarded. If the monkey released the bar before reaching the time criterion or touched the target in more than 1 s after bar release, it was not rewarded. In these cases, the trial continued until the end of its total duration, and the same trial was repeated until a rewarded movement was successfully completed. Trials on which animals did not react after cue onset or did not touch a target after bar release were excluded from the analyses. Both monkeys received training on the TET until they reached a stable level of performance corresponding to a criterion of at least 80% correctly performed trials.

We also tested the effects of intervals in the seconds range in a Pavlovian conditioning task (PCT) that excludes possible confounding factors that were linked to target reaching. In this condition, access to the working panel was prevented by closing the sliding door at the front of the restraining box and monkeys remained motionless with their arms relaxed in a natural position. Animals were then exposed to a visual stimulus (red light, 0.5 s duration), presented either on the left or right side, whose onset was initiated by the experimenter. This cue was followed by the delivery of reward after a fixed interval of 1.0 or 2.0 s, depending upon stimulus location. The PCT matched the TET in terms of location-interval combinations (left/short and right/long) and total trial duration (6 s). During PCT sessions, the licking movements of the monkeys were monitored using force transducers (strain gauges) attached to the tube delivering liquid. Signals from the strain gauge device were digitized at 500 Hz and stored into an analog file. In addition, we sometimes reversed the cue-interval associations of the PCT. By comparing TAN activity before and after reversal, we assessed whether activity changes following the cue onset were related to the ensuing interval duration independent of the cue location.

### Data analysis

Performance in the TET was assessed by measuring the time between the onset of the cue and bar release (movement onset time, MOT), for each time-interval combination. We examined MOT distributions to determine whether the spread of these distributions is proportional to the length of the interval being estimated, as reflected by the ratio between the dispersion and the mean of MOT (coefficient of variation, CV).

Neurons were determined to be task-responsive according to methods similar to those described previously (Marche et al., 2017). Briefly, a “sliding window” analysis was used to quantify the time course of changes in TAN activity after a specific task event. A test window of 100 ms duration was advanced in 10 ms increments across the trial, starting at cue onset, movement onset, or target contact. The average spike counts in the test window was compared, at each step, with that calculated during the 500 ms immediately preceding the onset of the cue (control period). The onset of a modulation was taken to be the beginning of the first of at least five consecutive steps showing a significant difference (Dunn’s test for multiple nonparametric comparisons, p < 0.01) as against the activity in the control period.

Quantitative analysis of spike rates was also performed for windows with fixed durations that were specifically determined for each monkey on the basis of the analysis of latency and duration of statistically significant decreases in activity (i.e., pauses) detected with the sliding window procedure. We set these time windows such that they included most of the pause onset and offset times. According to this analysis, time windows after the cue onset were 93-309 ms for both short- and long-interval trials, for monkey C, and 127-313 ms and 211-418 ms for short- and long-interval trials respectively, for monkey D. The windows determined by the average onset and offset times of pause responses to reward were 139-369 ms and 194–391 ms for both short- and long-interval trials, for monkey C and D, respectively. We then rated the magnitude of activity changes in each time window for each neuron and each time interval, irrespective of whether neurons were individually responsive or not. We set quantification windows in the same way for assessing the magnitude of TAN activity after the stimulus and reward in the PCT. The percentages of TANs showing pause responses in relation to the total number of neurons tested were calculated for each condition and differences in proportions of responding neurons among conditions were statistically assessed with the χ^2^ test. Differences in the magnitude of changes in activity were assessed with a Wilcoxon rank-sum test.

### Recording sites

The location of individual recorded neurons was confirmed histologically in both monkeys. Near the end of the experiments, small electrolytic lesions (20 μA for 30 s, cathodal current) were made at several points along selected electrode tracks. The monkey was deeply anesthetized with pentobarbital and perfused through the heart with 0.9% saline followed by 4% paraformaldehyde (pH 7.4 phosphate buffer). Frozen sections (40-μm thick) were made in the frontal plane and stained with cresyl violet. We then reconstructed the location of each recorded neuron according to the depth and coordinates of electrode penetrations based on the retrieved sites of marking lesions. In line with previous work, we used the AC as a structural boundary separating the striatum into motor and associative parts. Based on the reconstruction of recording sites, TANs were sampled between 5 mm anterior and 4 mm posterior to the AC, mainly over the medio-lateral extent of the putamen. Only a few penetrations were made in the most ventral part of the precommissural striatum.

## Acknowledgements

We thank Dr. L. Renaud for assistance with surgery, Dr. K. Marche for assistance in MATLAB programming, and Dr. M. Esclapez for help with histology. We also thank animal care personnel of the Mediterranean Primate Research Center.

## Data availability statement

All datasets generated for this study will be made available upon publication. To make our data openly accessible, we will consider sharing source data files and codes by using a public open repository.

---

**Figure 2-figure supplement 1.**
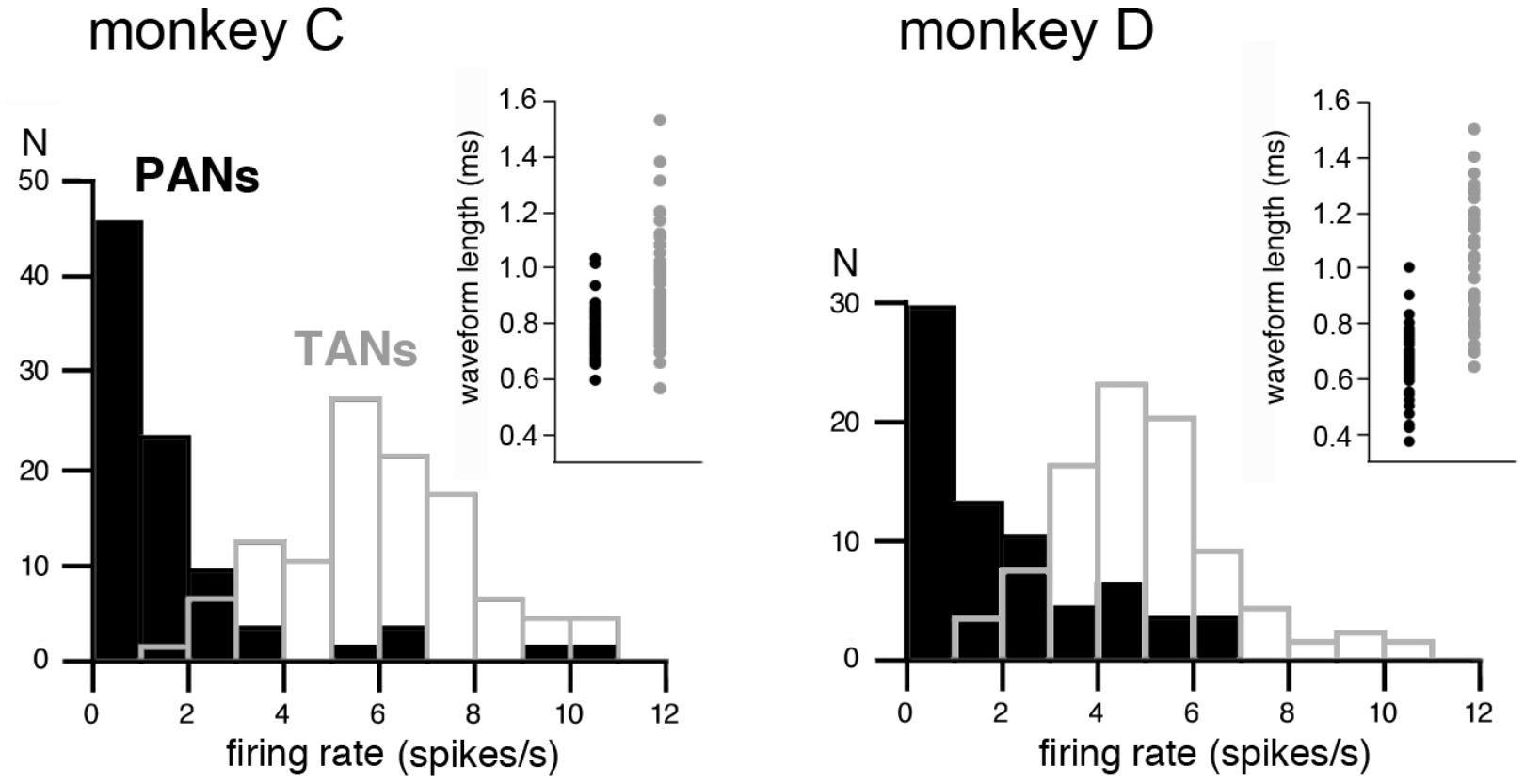
Electrophysiological characterization of TANs. Tonically active neurons (TANs), presumed cholinergic interneurons in the striatum, were identified based on previously established electrophysiological criteria, i.e., spontaneous firing rate and spike waveform width. Consistent with previous studies, the TANs had a mean baseline activity which is higher than that of the phasically active neurons (PANs), which correspond to striatal output neurons (Monkey C: TANs: n=103, 5.94 + 1.94 spikes/s; PANs: n=86, 1.47 + 1.91 spikes/s; Monkey D: TANs: n=86, 4.82 + 1.72 spikes/s; PANs: n=68, 1.83 + 1.92 spikes/s). Also, the mean spike waveform width of TANs was larger than that of the PANs (Monkey C: TANs: n=109, 0.90 + 0.17 ms; PANs: n=104, 0.76 + 0.08 ms; Monkey D: TANs: n=85, 1.08 + 0.22 spike/s; PANs: n=68, 0.65 + 0.11 ms). In addition, the stereotyped *pause* in activity in response to motivationally salient stimuli is a reliable indicator of TAN identity which clearly distinguishes them from PANs. We excluded from analysis a relatively small sample of the so-called fast spiking interneurons, presumed striatal GABAergic interneurons, characterized by having a higher firing frequency than TANs and a narrower spike waveform than PANs. Spontaneous activity was measured as the mean firing rate during the 500 ms before the onset of the timing cue. N indicates the number of neurons. Insets: distributions of average spike lengths. Each dot represents data from an individual neuron. The spike length of each neuron was defined as the time interval between the first negative and second positive peaks of the waveform.

**Figure 4-figure supplement 1.**
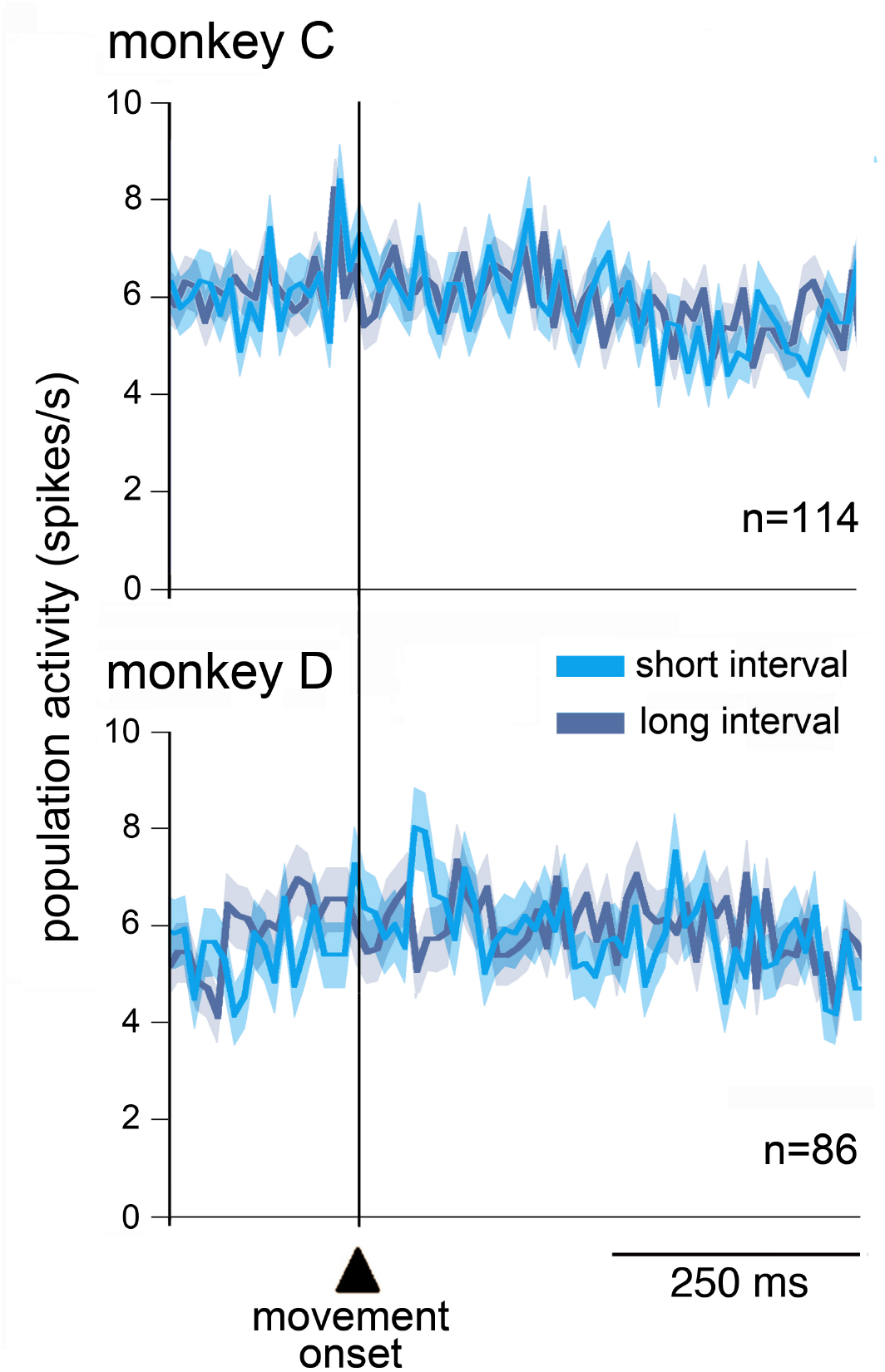
Lack of changes in TAN activity around self-timed movements. Population activity of TANs aligned on the onset of movements marked by the vertical line. At a population level, TANs did not exhibit clear activity modulation around the movement onset, regardless of the time interval. Same conventions as in Fig. 2D.

**Figure 5-figure supplement 1.**
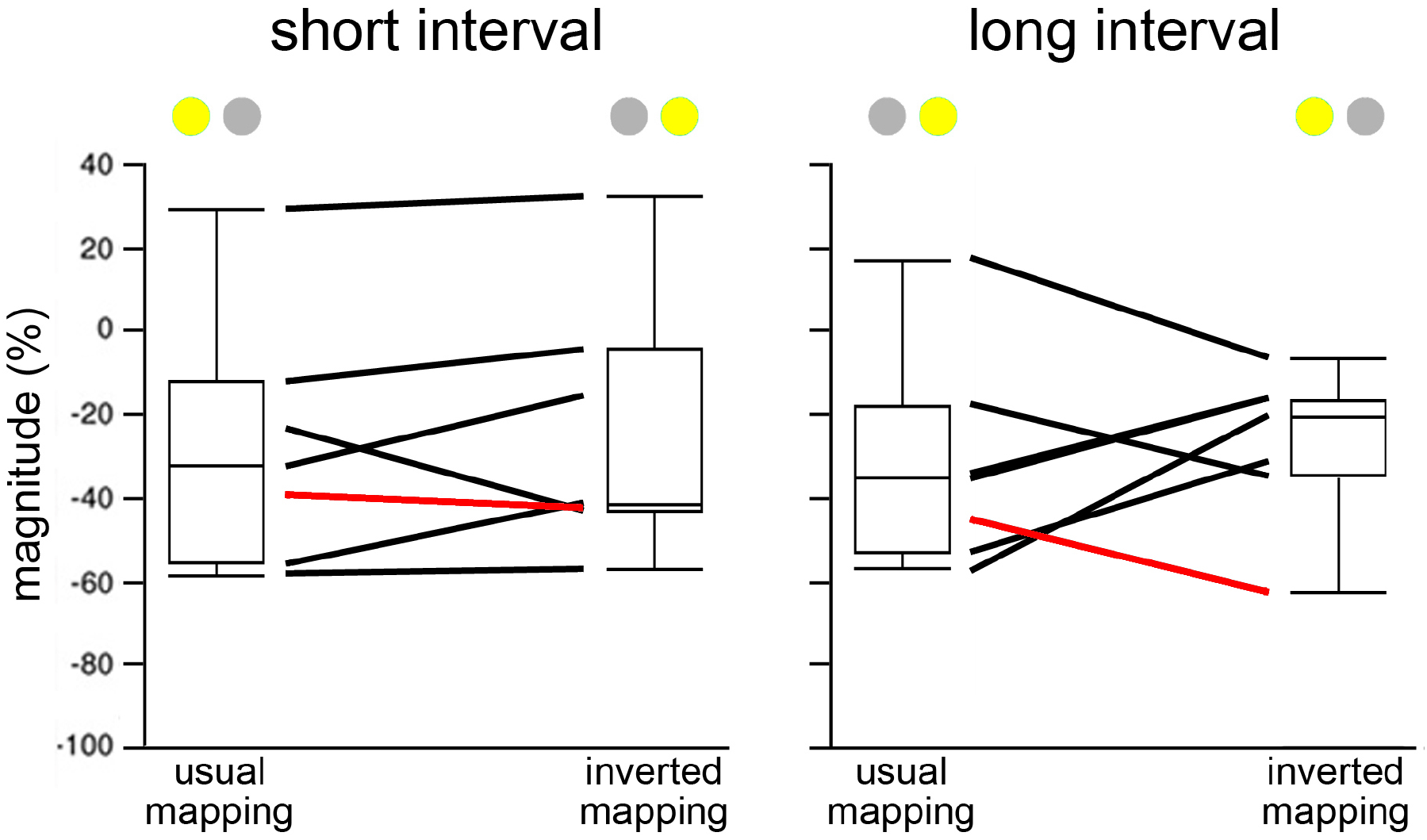
Influence of changing the cue-interval mapping in the Pavlovian protocol. Box plots of the magnitude of changes in TAN activity after the onset of the cue, separately for short and long intervals. Boxes indicate 25–75 percentile ranges of the distributions, lines through boxes indicate medians and vertical lines indicate tails of distributions. Red lines correspond to the example TAN illustrated in Fig. 5B. We recorded a sample of seven TANs in the PCT to test whether TAN responses to the cue are attributable to the time interval or the location of the cue. After carrying out a block of trials with the usual assignment of cues for the two time intervals (usual mapping), we reversed the assignments of the spatial cues for the short and long interval (inverted mapping). The activity across this sample of TANs was not altered when the cue-interval mapping was reversed, suggesting that it was not influenced by the location of the cue.

## References

Apaydin N, Üstün S, Kale EH, Çelikağ I, Özgüven HD, Baskak B, Çiçek M. 2018. Neural mechanisms underlying time perception and reward anticipation. Frontiers in Human Neuroscience 12:115. DOI: 10.3389/fnhum.2018.00115

Aoki S, Liu AW, Zucca A, Zucca S, Wickens JR. 2015. Role of striatal cholinergic interneurons in set-shifting in the rat. The Journal of Neuroscience 35:9424–9431. DOI: 10.1523/JNEUROSCI.0490-15.2015

Aosaki T, Tsubokawa H, Ishida A, Watanabe K, Graybiel AM, Kimura M. 1994. Responses of tonically active neurons in the primate’s striatum undergo systematic changes during behavioral sensorimotor conditioning. The Journal of Neuroscience 14:3969–3984. DOI: 10.1523/JNEUROSCI.14-06-03969.1994

Apicella P. 2007. Leading tonically active neurons of the striatum from reward detection to context recognition. Trends in Neuroscience 30:299–306. DOI: 10.1016/j.tins.2007.03.011

Apicella P. 2017. The role of the intrinsic cholinergic system of the striatum: What have we learned from TAN recordings in behaving animals? Neuroscience 360:81–94. DOI: 10.1016/j.neuroscience.2017.07.060

Apicella P, Legallet E, Trouche E. 1997. Responses of tonically discharging neurons in the monkey striatum to primary rewards delivered during different behavioral states. Experimental Brain Research 116:456–466. DOI: 10.1007/pl00005773

Apicella P, Ravel S, Legallet E. 2006. A possible role for tonically active neurons of the primate striatum in learning about temporal relationships among salient stimuli. In: E. Bezard (Ed) Recent Breakthroughs in Basal Ganglia Research. Nova Science Publishers, New York. pp. 55–63.

Apicella P, Deffains M, Ravel S, Legallet E. 2009. Tonically active neurons in the striatum differentiate between delivery and omission of expected reward in a probabilistic task context. The European Journal of Neuroscience 30:515–526. DOI: 10.1111/j.1460-9568.2009.06872

Apicella P, Ravel S, Deffains M, Legallet E. 2011. The role of striatal tonically active neurons in reward prediction error signaling during instrumental task performance. The Journal of Neuroscience 31:1507–1515. DOI: 10.1523/JNEUROSCI.4880-10.2011

Avlar B, Kahn JB, Jensen G, Kandel ER, Simpson EH, Balsam PD. 2015. Improving temporal cognition by enhancing motivation. Behavioral Neuroscience 129:576–588. DOI: 10.1037/bne0000083

Bakhurin KI, Mac V, Golshani P, Masmanidis SC. 2016. Temporal correlations among functionally specialized striatal neural ensembles in reward conditioned mice. Journal of Neurophysiology 115:1521–1532. DOI: 10.1152/jn.01037.2015

Bakhurin KI, Goudar V, Shobe JL, Claar LD, Buonomano DV, Masmanidis SC. 2017. Differential encoding of time by prefrontal and striatal network dynamics. The Journal of Neuroscience 37:854–870. DOI: 10.1523/JNEUROSCI.1789-16.2016

Balci F. 2014. Interval timing, dopamine, and motivation. Timing and time perception 2:379–410. DOI: 10.1163/22134468-00002035

Bradfield LA, Bertran-Gonzalez J, Chieng B, Balleine BW. 2013. The thalamostriatal pathway and cholinergic control of goal-directed action: interlacing new with existing learning in the striatum. Neuron 79:153–166. DOI: 10.1016/j.neuron.2013.04.039

Brown HD, Baker PM, Ragozzino ME. 2010. The parafascicular thalamic nucleus concomitantly influences behavioral flexibility and dorsomedial striatal acetylcholine output in rats. The Journal of Neuroscience 30:14390–14398. DOI: 10.1523/JNEUROSCI.2167-10.2010

Buhusi CV, Meck WH. 2005. What makes us tick? Functional and neural mechanisms of interval timing. Nature Reviews Neuroscience 6:755. DOI: 10.1038/nrn1764

Cachope R, Mateo Y, Mathur BN, Irving J, Wang H-L, Morales M, Lovinger DM, Cheer JF. 2012. Selective activation of cholinergic interneurons enhances accumbal phasic dopamine release: setting the tone for reward processing. Cell Report 2:33–41. DOI: 10.1016/j.celrep.2012.05.011

Cai Y, Ford CP. 2018. Dopamine cells differentially regulate striatal cholinergic transmission across regions through corelease of dopamine and glutamate. Cell Report 25:3148–3157. DOI: 10.1016/j.celrep.2018.11.053

Chiba A, Oshio KI, Inase M. 2015. Neuronal representation of duration discrimination in the monkey striatum. Physiological Reports 3:e12283. doi: 10.14814/phy2.12283

Coull JT, Cheng R-K., Meck WH. 2011. Neuroanatomical and neurochemical substrates of timing. Neuropsychopharmacology 36:3–25. DOI: 10.1038/npp.2010.113

Daniels CW, Sanabria F. 2017. Interval timing under a behavioral microscope: Dissociating motivational and timing processes in fixed-interval performance. Learning and Behavior 45:29–48. DOI: 10.3758/s13420-016-0234-1

Daw ND, Courville AC, Tourtezky DS. 2006. Representation and timing in theories of the dopamine system. Neural Computation 18:1637–1677. DOI: 10.1162/neco.2006.18.7.1637

Ding JB, Guzman JN, Peterson JD, Goldberg JA, Surmeier DJ. 2010. Thalamic gating of corticostriatal signaling by cholinergic interneurons. Neuron 67:294–307. DOI: 10.1016/j.neuron.2010.06.017

Fung BJ, Sutlief E, Shuler MGH. 2021. Dopamine and the interdependency of time perception and reward. Neuroscience Biobehavioral Review 125:380–391. DOI: 10.1016/j.neubiorev.2021.02.030

Gable PA, Poole BD. 2012. Time flies when you’re having approach-motivated fun: effects of motivational intensity on time perception. Psychological Science 23:879–86. DOI: 10.1177/0956797611435817.

Galtress T, Marshall AT, Kirkpatrick K. 2012. Motivation and timing: Clues for modeling the reward system. Behavioural Processes 90:142–153. DOI: 10.1016/j.beproc.2012.02.014

Gershman SJ, Moustafa AA, Ludvig EA. 2014. Time representation in reinforcement learning models of the basal ganglia. Frontiers in Computational Neuroscience 7:194. DOI: 10.3389/fncom.2013.00194

Gibbon J. 1977. Scalar expectancy theory and Weber’s law in animal timing. Psychological Review 84:279–325. DOI: 10.1037/0033-295X.84.3.279

Gouvêa TS, Monteiro T, Motiwala A, Soares S, Machens C, Paton JJ. 2015. Striatal dynamics explain duration judgments. eLife 4:e11386. DOI: 10.7554/eLife.11386

Graziola F, Pellorca C, Di Criscio L, Vigevano F, Curatolo P, Capuano A. 2020. Impaired motor timing in Tourette syndrome: results from a case–control study in children. Frontiers in Neurology 11:552701. DOI: 10.3389/fneur.2020.552701

Harrington DL, Boyd LA, Mayer AR, Sheltraw DM, Lee RR, Huang M, Rao SM. 2004. Neural representation of interval encoding and decision making. Cognitive Brain Research 21:193–205. DOI: 10.1016/j.cogbrainres.2004.01.010

Howe M, Ridouh I, Allegra Mascaro AL, Larios A, Azcorra M, Dombeck DA. 2019. Coordination of rapid cholinergic and dopaminergic signaling in striatum during spontaneous movement. eLife 8:e44903. DOI: 10.7554/eLife.44903

Jin DZ, Fujii N, Graybiel AM. 2009. Neural representation of time in cortico-basal ganglia circuits. PNAS 106:19156–19161. DOI: 10.1073/pnas.0909881106

Kataoka Y, Kalanithi PSA, Grantz H, Schwartz ML, Saper C, Leckman JF, Vaccarino FM. 2010. Decreased number of parvalbumin and cholinergic interneurons in the striatum of individuals with Tourette syndrome. The Journal of Comparative Neurology 518:277–291. DOI: 10.1002/cne.22206

Kimura M. 1992. Behavioral modulation of sensory responses of primate putamen neurons. Brain Research 578:204–214. DOI: 10.1016/0006-8993(92)90249-9

Kimura M, Matsumoto N, Okahashi K, Ueda Y, Satoh T, Minamimoto T, Sakamoto M, Yamada H. 2003. Goal-directed, serial and synchronous activation of neurons in the primate striatum. Neuroreport 14:799–802. DOI: 10.1097/00001756-200305060-00004

Kirkpatrick K. 2014. Interactions of timing and prediction error learning. Behavioural Processes 101:135–145. DOI: 10.1016/j.beproc.2013.08.005

Lee IH, Seitz AR, Assad JA. 2006. Activity of tonically active neurons in the monkey putamen during initiation and withholding of movement. Journal of Neurophysiology 9:2391–2403. DOI: 10.1152/jn.01053.2005

Marche K, Martel AC, Apicella P. 2017. Differences between dorsal and ventral striatum in the sensitivity of tonically active neurons to rewarding events. Frontiers in Systems Neuroscience 11:52. DOI: 10.3389/fnsys.2017.00052.

Martel AC, Apicella P. 2021. Temporal processing in the striatum: Interplay between midbrain dopamine neurons and striatal cholinergic interneurons. European Journal of Neuroscience 53:2090–2099. DOI: 10.1111/ejn.14741.

Matell MS, Meck WH. 2000. Neuropsychological mechanisms of interval timing behavior. BioEssays 22:94–103. DOI: 10.1002/(SICI)1521-1878(200001)22:1<94::AID-BIES14>3.0.CO;2-E

Matell MS, Meck WH. 2004. Cortico-striatal circuits and interval timing: coincidence detection of oscillatory processes. Cognitive Brain Research 21:139–170. DOI: 10.1016/j.cogbrainres.2004.06.012

Matell MS, Meck WH, Nicolelis MAL. 2003. Interval timing and the encoding of signal duration by ensembles of cortical and striatal neurons. Behavioral Neuroscience 117:760–773. DOI: 10.1037/0735-7044.117.4.760

Matsumoto N, Minamimoto T, Graybiel AM, Kimura M. 2001. Neurons in the thalamic CM-Pf complex supply striatal neuons with information about behaviorally significant sensory events. Journal of Neurophysiology 85:60–976. DOI: 10.1152/jn.2001.85.2.960

Meck WH. 1996. Neuropharmacology of timing and time perception. Cognitive Brain Research 3:227–242. DOI: 10.1016/0926-6410(96)00009-2

Meck WH. 2006. Neuroanatomical localization of an internal clock: A functional link between mesolimbic, nigrostriatal, and mesocortical dopaminergic systems. Brain Research 1109:93–107. DOI: 10.1016/j.brainres.2006.06.031

Mello GB, Soares S, Paton JJ. 2015. A scalable population code for time in the striatum. Current Biology 25:1113–2112. DOI: 10.1016/j.cub.2015.02.036

Merchant H, Harrington DL, Meck WH. 2013. Neural basis of the perception and estimation of time. Annual Review of Neuroscience 36:313–336. DOI: 10.1146/annurev-neuro-062012-170349

Mikhael JG, Gershman SJ. 2019. Adapting the flow of time with dopamine. Journal of Neurophysiology 121:1748–1760. DOI: 10.1152/jn.00817.2018

Morris G, Arkadir D, Nevet A, Vaadia E, Bergman H. 2004. Coincident but distinct messages of midbrain dopamine and striatal tonically active neurons. Neuron 43:133–143. DOI: 10.1016/j.neuron.2004.06.012

Noreika V, Falter CM, Rubia K. 2013. Timing deficits in attention-deficit/hyperactivity disorder (ADHD): evidence from neurocognitive and neuroimaging studies. Neuropsychologia 51:235–266. DOI: 10.1016/j.neuropsychologia.2012.09.036.

Parker KL, Lamichhane D, Caetano MS, Narayanan NS. 2013. Executive dysfunction in Parkinson’s disease and timing deficits. Frontiers in Integrative Neuroscience 7. DOI: 10.3389/fnint.2013.00075

Pastor MA, Artieda J, Jahanshahi M, Obeso JA. 1992. Time estimation and reproduction is abnormal in Parkinson’s disease. Brain 115:211–225. DOI: 10.1093/brain/115.1.211

Paton JJ, Buonomano DV. 2018. The neural basis of timing: distributed mechanisms for diverse functions. Neuron 98:687–705. DOI: 10.1016/j.neuron.2018.03.045

Paxinos G, Huang X, Petrides M, Toga A. 2008. The Rhesus Monkey Brain in Stereotaxic Coordinates. 2nd edn. San Diego: Academic Press.

Petter EA, Gershman SJ, Meck WH. 2018. Integrating models of interval timing and reinforcement learning. Trends in Cognitive Sciences 22:911–922. DOI: 10.1016/j.tics.2018.08.004

Ravel S, Sardo P, Legallet E, Apicella P. 2001. Reward unpredictability inside and outside of a task context as a determinant of the responses of tonically active neurons in the monkey striatum. The Journal of Neuroscience 21:5730–5739. DOI: 10.1523/JNEUROSCI.21-15-05730.2001

Ravel S, Sardo P, Legallet E, Apicella P. 2006. Influence of spatial information on responses of tonically active neurons in the monkey striatum. Journal of Neurophysiology 95:2975–2986. DOI: 10.1152/jn.01113.2005

Raz A, Feingold A, Zelanskaya V, Vaadia E, Bergman H. 1996. Neuronal synchronization of tonically active neurons in the striatum of normal and parkinsonian primates. Journal of Neurophysiology 76:2083–2088. DOI: 10.1152/jn.1996.76.3.2083

Rubia K, Halari R, Christakou A, Taylor E. 2009. Impulsiveness as a timing disturbance: neurocognitive abnormalities in attention-deficit hyperactivity disorder during temporal processes and normalization with methylphenidate. Philosophical transactions of the royal society B 364:1919–1931. DOI: 10.1098/rstb.2009.0014

Sardo P, Ravel S, Legallet E, Apicella P. 2000. Influence of the predicted time of stimuli eliciting movements on responses of tonically active neurons in the monkey striatum. European Journal of Neuroscience 12:1801–1816. DOI: 10.1046/j.1460-9568.2000.00068.x

Schüller CB, Wagner BJ, Schüller T, Baldermann JC, Huys D, Kerner Auch Koerner J, Niessen E, Münchau A, Brandt V, Peters J, Kuhn J. 2021. Temporal discounting in adolescents and adults with Tourette syndrome. PLoS 16:e0253620. DOI: 10.1371/journal.pone.0253620

Schulz JM, Reynolds JN. 2013. Pause and rebound: sensory control of cholinergic signaling in the striatum. Trends in Neuroscience 36:41–50. DOI: 10.1016/j.tins.2012.09.006.

Shimo Y, Hikosaka O. 2001. Role of tonically active neurons in primate caudate in reward-oriented saccadic eye movement. The Journal of Neuroscience 21:7804–7814. DOI: 10.1523/JNEUROSCI.21-19-07804.2001

Soares S, Atallah BV, Paton JJ. 2016. Midbrain dopamine neurons control judgment of time. Science 354:1273–1277. DOI: 10.1126/science.aah5234

Stalnaker TA, Berg B, Aujla N, Schoenbaum G. 2016. Cholinergic interneurons use orbitofrontal input to track beliefs about current State. The Journal of Neuroscience 36:6242–6257. DOI: 10.1523/JNEUROSCI.0157-16.2016

Stocco A. 2012. Acetylcholine-based entropy in response selection: a model of how striatal interneurons modulate exploration, exploitation, and response variability in decision-making. Frontiers in Neuroscience 6:18. DOI: 10.3389/fnins.2012.00018

Threlfell S, Cragg SJ. 2011. Dopamine signaling in dorsal versus ventral striatum: the dynamic role of cholinergic interneurons. Frontiers in Systems Neuroscience 5. DOI: 10.3389/fnsys.2011.00011

Threlfell S, Lalic T, Platt NJ, Jennings KA, Deisseroth K, Cragg SJ. 2012. Striatal dopamine release is triggered by synchronized activity in cholinergic interneurons. Neuron 75:58–64. DOI: 10.1016/j.neuron.2012.04.038

Tomasi D, Wang GJ, Studentsova Y, Volkow ND. 2015. Dissecting neural responses to temporal prediction, attention, and memory: effects of reward learning and interoception on time perception. Cerebral Cortex 25:3856–3867. DOI: 10.1093/cercor/bhu269

Vicario CM, Martino D, Spata F, Defazio G, Giacchè R, Martino V, Rappo G, Pepi AM, Silvestri PR, Cardona F. 2010. Time processing in children with Tourette’s syndrome. Brain and Cognition 73:28–34. DOI: 10.1016/j.bandc.2010.01.008

Wang J, Narain D, Hosseini EA, Jazayeri M. 2018. Flexible timing by temporal scaling of cortical responses. Nature Neuroscience 21:102–110. DOI: 10.1038/s41593-017-0028-6

Yamada H, Matsumoto N, Kimura M. 2004. Tonically active neurons in the primate caudate nucleus and putamen differentially encode instructed motivational outcomes of action. The Journal of Neuroscience 24:3500–3510. DOI: 10.1523/JNEUROSCI.0068-04.2004

Zhou F-M, Wilson CJ, Dani JA. 2002. Cholinergic interneuron characteristics and nicotinic properties in the striatum. Journal of Neurobiology 53:590–605. DOI: 10.1002/neu.10150

Zhou S, Masmanidis SC, Buonomano DV. 2020. Neural sequences as an optimal dynamical regime for the readout of time. Neuron 108:651–658.e5. DOI: 10.1016/j.neuron.2020.08.020.

